# Variation in TAF1 expression in female carrier induced pluripotent stem cells and human brain ontogeny has implications for adult neostriatum vulnerability in X-linked Dystonia Parkinsonism

**DOI:** 10.1101/2022.03.26.485862

**Authors:** Laura D’Ignazio, Ricardo S. Jacomini, Bareera Qamar, Kynon J.M. Benjamin, Ria Arora, Tomoyo Sawada, Taylor A. Evans, Kenneth E. Diffenderfer, Aimee R. Pankonin, William T. Hendriks, Thomas M Hyde, Joel E Kleinman, Daniel R Weinberger, D. Cristopher Bragg, Apua C.M. Paquola, Jennifer A. Erwin

## Abstract

X-linked Dystonia-Parkinsonism (XDP) is an inherited, X-linked, adult-onset movement disorder characterized by degeneration in the neostriatum. No therapeutics alter disease progression. The mechanisms underlying regional differences in degeneration and age of onset are unknown. Developing therapeutics that target XDP-related mechanisms requires a deeper understanding of how XDP-relevant features vary in health and disease. XDP is due, in part, to either a partial loss of *TAF1* function and/or a SVA-driven pathological gain of function. A disease-specific SINE-VNTR-Alu (SVA) retrotransposon insertion occurs within intron 32 of *TAF1*, a subunit of TFIID involved in transcription initiation. While all XDP males are usually clinically affected, females are heterozygous carriers generally not manifesting the full syndrome. As a resource for disease modeling, we characterized eight iPSC lines from XDP female carrier individuals, and identified isogenic lines where one clonal iPSC line expressed the wild-type X, and the two other clonal iPSC lines expressed the XDP haplotype. Furthermore, we characterized XDP-relevant transcript expression variation in humans, and found that SVA-F expression decreases slightly after 30 years of age in the neurotypical human brain and that *TAF1* is modestly decreased in the majority of female samples. Uniquely in the caudate nucleus, *TAF1* expression is not sexually dymorphic and decreased after 15 years of age. These findings indicate that regional-, age- and sex-specific mechanisms regulate *TAF1*, highlighting the importance of disease-relevant models and postmortem tissue analysis. We propose that the decreased *TAF1* expression in the adult caudate may synergize with the XDP-specific partial loss of *TAF1* function in patients, thereby passing a minimum threshold of *TAF1* function, and triggering degeneration in the neostriatum.

**Significance Statement:** XDP is an inherited, X-linked, adult-onset movement disorder characterized by degeneration in the neostriatum. No therapeutics alter disease progression. Developing therapeutics requires a deeper understanding of how XDP-relevant features vary in health and disease. XDP is possibly due to a partial loss of *TAF1* function. While all XDP males are usually affected, females are heterozygous carriers generally not manifesting the full syndrome. As a resource for disease modeling, we characterized eight stem cell lines from XDP female carrier individuals. Furthermore, we found that, uniquely in the caudate nucleus, *TAF1* expression decreases after adolescence in healthy humans. We hypothesize that the decrease of *TAF1* after adolescence in human caudate, in general, may underlie the vulnerability of the adult neostriatum in XDP.

## Introduction

X-linked Dystonia-Parkinsonism (XDP) is a rare movement disorder, predominantly affecting males descending from the island of Panay, in the Philippines (OMIM #3142590). Disease onset typically occurs within the third or fourth decade of life, and clinical manifestations consist of hyperkinetic movements and focal dystonia shifting toward parkinsonian features in a temporal evolution spanning 5-10 years (Lee et al., 2011). Treatment is limited to symptomatic management, including oral medications, botulinum toxin, and pallidal deep brain stimulation (Bragg et al., 2019).

Neuropathological analyses of a few XDP postmortem brains demonstrated progressive neuronal loss within the neostriatum, particularly affecting medium spiny neurons (MSNs) (Goto et al., 2013, 2005; Hanssen et al., 2018). Neuroimaging has detected other neostriatal and extra-neostriatal abnormalities (Arasaratnam et al., 2021; Blood et al., 2018; Brüggemann et al., 2016; Hanssen et al., 2018; Walter et al., 2017). Similar to many neurodegenerative disorders, the mechanisms underlying XDP neurodegeneration and the differential vulnerability to pathology are unknown.

XDP is a Mendelian disorder and thirteen shared variants have been detected in all XDP probands: 11 single disease-specific nucleotide changes (Aneichyk et al., 2018; Nolte et al., 2003), a 48 bp deletion (Nolte et al., 2003), and an ∼2.6 kb SVA-F type (short interspersed nuclear elements (SINE) - variable number of tandem repeats (VNTR) - Alu) retrotransposon insertion (Makino et al., 2007). All variants occur in intergenic or intronic regions within the *TAF1* gene (TATA-binding protein (TBP)-associated factor 1), encoding a core component of the TFIID complex that coordinates RNA polymerase II (RNA pol II) promoter initiation (Domingo et al., 2015).

XDP patient samples have decreased expression of the canonical full-length *TAF1* (*cTAF1*) transcript in dividing cells (Aneichyk et al., 2018; Rakovic et al., 2018), patient blood RNA (Al Ali et al., 2021), and a 2D cellular model of MSNs (Rakovic et al., 2018). The *cTAF1* decreases are caused by the SVA, which induces alternative splicing and intron retention (Aneichyk et al., 2018). In XDP cells, one of the most abundant aberrant transcripts caused by the pathogenic SVA is *TAF1-32i,* composed of canonical exon 32, spliced to a cryptic exon in intron 32, that terminates 5’ to the SVA insertion site (Aneichyk et al., 2018).

SVAs are non-coding hominid-specific human retrotransposons. There are roughly 2700 SVA copies per human genome, with the SVA-F being the evolutionary youngest (Hancks and Kazazian, 2010). SVA elements are regulated by protein coding genes (Savage et al. 2013), but the dynamics of SVA expression during human brain lifespan have not been well elucidated.

XDP is an X-linked recessive disorder, and the vast majority of affected cases are hemizygous males. Heterozygous carrier females typically do not manifest the full syndrome, and, to date, only 14 females exhibited clinical signs resembling the full syndrome compared to more than 500 affected hemizygous males (Domingo et al., 2014; Evidente et al., 2004). Female carriers without the full syndrome do demonstrate some pathology, including decreased caudate and putamen volumes compared to healthy controls (Blood et al., 2018). Understanding how XDP manifests in female carriers can reveal information about disease pathophysiology.

As a consequence of the dosage compensation mechanism X-chromosome inactivation (XCI), females transcriptionally silence most genes, including *TAF1* (Oliva et al., 2020), on one X chromosome (Patrat et al., 2020). During human embryo implantation, one X chromosome in each cell is randomly epigenetically silenced, with stable inheritance of the epigenetically silenced inactivated X chromosome in subsequent daughter cells. Therefore, tissues of heterozygous female XDP carriers are a mosaic of two cell populations, with either the wild-type or mutant X chromosome randomly inactivated per cell (Migeon, 2020).

Induced pluripotent stem cell (iPSC) models are a powerful human cellular system that can elucidate mechanisms of disease without the limitations of postmortem tissue access and end-state disease confounds. However, female iPSC have known X chromosome inactivation abnormalities (Geens and Chuva De Sousa Lopes, 2017). Therefore, here we perform extensive characterization of eight human induced pluripotent stem cell (iPSC) lines reprogrammed from fibroblasts derived from three XDP heterozygous female carriers (Aneichyk et al., 2018; Ito et al., 2016) to serve as a valuable tool to study XDP pathogenesis *in vitro* upon differentiation into specific neuronal models.

Furthermore, it is not yet known the natural level of *TAF1* and SVA expression variation in human. We hypothesize that age and tissue specific variation in *TAF1* and SVA expression may underlie the age and tissue-specific degeneration in XDP. Therefore, we performed extensive transcriptomic analyses of stem cell models and postmortem neurotypical brain regions to characterize TAF1 and SVA expression in relevant human tissue and cells.

## Materials and methods

### Cell culture of iPSC lines

Derivation and reprogramming from skin fibroblasts, as well as the initial characterization of iPSCs derived from a male patient with XDP, a male unaffected relative (control), and three XDP female carriers have previously been reported elsewhere (Aneichyk et al., 2018; Ito et al., 2016). The full list of iPSCs used in our study is reported in **Extended Data Table 1-1**, including the XDP isogenic SVA-deleted iPSC line (ΔSVA-XDP), previously generated through CRISPR/Cas9 genome editing (Aneichyk et al., 2018). All iPSCs were expanded and grown on Cultrex Growth Factor Reduced BME (Bio-Techne) in mTeSR or mTeSR Plus medium (Stemcell). The iPSCs were passaged every 4-6 days with Versene solution (Gibco) and split 1:8 to 1:10 each time. Cells were routinely tested for mycoplasma contamination using a MycoAlert Mycoplasma Detection Kit (Lonza).

### Ethics approval and consent

Performance of all experiments in this study strictly followed ethical guidelines, and any necessary Institutional Review Board (IRB) and/or ethics committee approvals were obtained. The cell lines used comprised de-identified induced pluripotent stem cells, which were derived previously (not specifically for the purpose of this study) from dermal punch fibroblasts at Massachusetts General Hospital (MGH) (Boston, MA). These specimens, sample collection, and derivation, have been approved by MGH IRB and were de-identified by MGH. These samples were transferred to the Lieber Institute for Brain Development (LIBD) under a Material Transfer Agreement (MTA). Later, the Western IRB granted an exemption under 45 CFR 46.101(b)(4), because our study involved the use of retrospective data that does not contain patient identifying information. We followed ethics guidelines set forth by LIBD and the Maryland Stem Cell Research Act.

Human postmortem brain tissues used for this study, and included in the BrainSeq Consortium, were collected at several sites. Most adult postmortem human brain samples were provided by the Clinical Brain Disorders Branch of the National Institute of Mental Health (NIMH) in collaboration with the Northern Virginia and District of Columbia Medical Examiners’ Offices, following a protocol approved by the IRB of the National Institutes of Health (NIH). Samples were transferred to the LIBD under a MTA with the NIMH. Additional samples were collected at the LIBD according to a protocol approved by the IRB of the State of Maryland Department of Health and Mental Hygiene and the Western IRB. Child and adolescent brain tissue samples were provided by the National Institute of Child Health and Human Development Brain and Tissue Bank for Developmental Disorders via MTAs approved by IRB of the University of Maryland. Audiotaped informed consent to study brain tissue was obtained from the legal next-of-kin on every case collected at NIMH and LIBD. Further details about donation process and specimens handling have been described previously (Lipska et al., 2006). Neurotypical postmortem brain samples used in this study were non-psychiatric non-neurological controls with no known history of significant psychiatric or neurological illnesses, including substance abuse. All necessary participant consent has been obtained and the appropriate institutional forms have been archived.

### RNA scope

An *in situ* hybridization assay was performed with RNAscope technology using the RNAscope Fluorescent Multiplex Kit V2 (ACD) according to the manufacturer’s instructions, and followed recommendations for cultured adherent cell preparations. Briefly, iPSCs were cultured in Cultrex-coated 4-well chambered coverslips (ibidi), and when 70-80% confluent, fixed with a 10% neutral buffered formalin solution (Sigma-Aldrich) for 30 min at room temperature. Then, iPSCs were series dehydrated in ethanol, pretreated with hydrogen peroxide for 10 min at room temperature, and treated with protease III diluted 1:15 in 1X PBS for 10 min. Cells were incubated with probes for XIST (#311231-C1, ACD), XACT (#471031-C2, ACD), HUWE1 (#526691-C2, ACD) and DUSP6 (#405361-C3, ACD), and stored overnight at 4°C in 4x SSC (saline-sodium citrate) buffer. The following day, probes were fluorescently labeled with Opal Dyes (Perkin Elmer) diluted in the TSA buffer. Specifically, Opal520 diluted 1:500 was assigned to HUWE1 and XACT; Opal570 diluted 1:500 was assigned to XIST; Opal690 diluted 1:500 was assigned to DUSP6. To label nuclei, we used DAPI (4′,6-diamidino-2-phenylindole). Imaging was performed using a laser confocal microscope (LSM700, Zeiss) and a 63X objective lens.

### Immunofluorescence staining and microscopy

For immunofluorescence staining, iPSCs were cultured in Cultrex-coated imaging dishes with a polymer coverslip bottom (ibidi). When colonies were 80-90% confluent, iPSCs were fixed with 4% PFA (Sigma) at room temperature for 15 min, permeabilized and blocked with 0.3% Triton X-100 (Sigma) and 10% normal horse serum (Thermo Fisher) in D-PBS (Gibco) for 30 min at room temperature. Cells were then stained overnight at 4°C, with two primary antibodies: goat anti-NANOG (1:200, #AF1997, R&D Systems) and mouse anti-TRA-1-60 (1:200, #MAB4360, Millipore Sigma), both diluted in 5% normal horse serum/0.01% Tween-20/D-PBS. The following day, iPSCs were incubated with donkey anti-goat IgG-AF488 conjugated (1:250, #705-545-147, Jackson ImmunoResearch), and donkey anti-mouse IgG-Cy3 conjugated (1:250, #715-165-151, Jackson ImmunoResearch), diluted in 0.01% Tween-20/D-PBS for 2 h at room temperature. Cell nuclei were labeled with Hoechst 33342 (1:10000, Invitrogen) for 10 min. Stained cells were imaged with laser confocal microscopy (LSM700, Zeiss) and processed with Zen software (Zeiss) and Adobe Photoshop CS4 (Adobe Systems, San Jose, CA).

### Genotyping and genome stability analysis

Genomic DNA was isolated from iPSCs with a DNeasy Blood and Tissue kit (Qiagen) as recommended by the manufacturer. Genotyping array analysis was performed by Psomagen with an Infinium Omni 2.5-8 kit (Illumina). To detect SVA, long-range PCR was performed following the method previously described by (Ito et al., 2016). The PCR amplicons were resolved by gel electrophoresis to identify a 3229 bp product (corresponding to the ∼2.6 kb SVA insertion), or a 599 bp product (without a SVA insertion).

### RNA extraction and Real Time quantitative PCR analysis

Total RNA was extracted from iPSCs with a Direct-zol Miniprep kit (Zymo Research). For reverse transcription, we used SuperScript IV VILO Master Mix (Thermo Fisher Scientific). The expression of XIST and TAF1 was analyzed via qPCR with a QuantiTect SYBR Green PCR kit (Qiagen) on a QuantStudio 3 (Applied Biosystems). The following 5 primer sets were used: XIST Forward: 5’-TTGCCCTACTAGCTCCTCGGAC-3’; XIST Reverse: 5’-TTCTCCAGATAGCTGGCAACC-3’; TAF1 Forward: 5’-AGAGTCGGGAGAGCTTTCTG-3’; TAF1 Reverse: 5’-CACAATCTCCTGGGCAGTCT-3’; GAPDH Forward: 5’-TGCACCACCAACTGCTTAGC-3’; GAPDH Reverse: 5’-GGCATGGACTGTGGTCATGAG-3’; HPRT Forward: 5’-TGACACTGGCAAAACAATGCA-3’; HPRT Reverse: 5’-GGTCCTTTTCACCAGCAAGCT-3’; GUSB Forward: 5’-GTCTGCGGCATTTTGTCGG-3’; GUSB Reverse: 5’-CACACGATGGCATAGGAATGG-3’.

To quantify intron 32 retention, we used a method previously described by (Aneichyk et al., 2018), as the authors generated a custom TaqMan primer/probe to detect the exon 32/intron 32 splice site (#AJWR28J, Thermo Fisher Scientific). When screening the XDP female carrier iPSC lines, 500 ng of RNA was reverse transcribed into cDNA using Superscript III First Strand Synthesis SuperMix (Thermo Fisher Scientific) with oligo(dT). The resulting cDNA was then further amplified through nested PCR using Phusion 2X Hot Start Flex MasterMix (NEB), as recommended by the manufacturer, with the following primers: TAF1-nested Forward: 5’-ACATCTCCAAGCACAAGTATCA-3’; TAF1-nested Reverse: 5’-GTAATGTACCAATATAAATTTCCTGGTTT-3’. The thermocycler conditions used for the nested PCR were: 98°C for 30 sec; 5 cycles of 98°C for 10 sec, 61°C for 30 sec, 72°C for 30 sec; 72°C for 5 min; and infinite hold at 4°C.

The nested PCR products were purified using a DNA clean/concentrator kit (Zymo Research) following the manufacturer’s instructions. Two microliters (∼25 ng/µl) of purified PCR products were used as template for subsequent qPCR performed on a QuantStudio 3 (Applied Biosystems) using TaqMan Fast Advanced Master Mix (Thermo Scientific), and the following conditions: 95°C for 20 sec; 40 cycles of 95°C for 1 sec, and 60°C for 20 sec. The GUSB probe/primer (#Hs009627_m1, Thermo Fisher Scientific) was used as a normalizer for gene expression analysis.

After qPCR analyses, for statistical purposes, one-way ANOVA followed by Dunnett’s test was used for comparing multiple conditions to a control condition only; one-way ANOVA followed by Tukey’s test was used for multiple pairwise condition comparisons. In all cases, p-values were determined as follows: * = p<0.05; ** = p<0.01 and *** = p<0.001.

### Allele-specific transcriptomic analysis

Total RNA was extracted from the eight XDP female carrier-derived iPSC lines, and libraries were prepared using the TruSeq Stranded Total RNA Gold kit to remove both cytoplasmic and mitochondrial rRNA. Paired-end 150 bp Illumina sequencing was performed at a median depth of 30M paired reads. Correction and filtering for technical errors (such as non unique alignments, low-quality bases, overlapping mates and duplicate reads), were carried out to remove potential sequencing errors and to minimize false-positive SNP identification. Reads were trimmed with Trimmomatics [0.39; (Bolger et al., 2014)]. To assess read quality before and after trimming, we performed FastQC (v0.11.8; https://github.com/s-andrews/FastQC). Then, we mapped the trimmed reads to GENCODE v32 (hg38/GRCh38) primary assembly using STAR aligner [2.7.3a; (Dobin et al., 2013)]. We examined alignment and additional read quality metrics with RSeQC [3.0.1; (L. Wang et al., 2012)].

To identify isogenic clonal pairs, single nucleotide variants (SNVs) were called from X chromosome-derived transcripts in all clones derived from the same XDP carrier individual, (33360, 33811, or 33110), using GATK best practices for SNPs and indel calling on RNA-Seq data. More specifically, allele-specific expression was analyzed following the GATK framework and GATK ASEReadCounter. Clonal pairs with the majority of SNVs unique to each clone were identified as isogenic pairs, as different X chromosomes were inactive in the two distinct lines. Conversely, clone pairs sharing most of SNVs were identified as iPSC lines with the same X chromosome inactivated.

### Global gene expression analysis

Gene counts were generated via featureCount [v1.5.3; (Liao et al., 2014)], a SubRead utility, for paired end, reversed stranded, reads in parallel (Tange, 2018), with four read filters: 1) uniquely mapped reads corresponding to a mapping quality of 60 for STAR BAM files; 2) reads aligned in proper pairs; 3) excluding chimeric reads; and 4) primary alignments. Raw counts were filtered for low expression by using filterByExpr from the edgeR Bioconductor package (McCarthy et al., 2012; Robinson et al., 2010), normalized for library size, followed by log2 counts per million (CPM) transformation.

### Differential expression analysis of the HipSci iPSC dataset

We downloaded FASTQ files associated with human iPSCs from (Kilpinen et al., 2017)) (n = 762; #PRJEB7388). Read quality was examined with FastQC prior to alignment to GENCODE v32 with HISAT2 [v2.1.0; (Kim et al., 2015)] and Salmon [v1.2.1; (Patro et al., 2017)]. For HISAT2 aligned reads, sorted BAM files were generated and indexed using SAMtools (v1.9). We generated gene counts with featureCount for paired end, reversed stranded reads similar to XDP gene counts using three read filters: 1) reads aligned in proper pairs; 2) excluding chimeric reads; and 3) primary alignments.

We selected samples based on two inclusion criteria: 1) British ethnicity of origin and 2) Caucasian ethnic group. This resulted in a total of 746 hiPSCs, comprising 438 females and 308 males, from 323 unique cell lines, with an age range from 25 to 79.

We performed quality control in two steps. In the first, we removed outliers by expression. To visualize outliers, we applied UMAP (uniform manifold approximation and projection) dimensional reduction to normalize Salmon-generated TPM (transcripts per million) log2 transformed expression. This resulted in two separated clusters, for which we dropped the smaller cluster, resulting in 662 hiPSCs. In the second quality control step, we performed RSeQC and aggregated the results of HISAT2 and FastQC using MultiQC (Ewels et al., 2016). We determined outliers based on read quality using principal component analysis (PCA) and centroid analysis. Specifically, we applied PCA to read quality metrics aggregated from MultiQC. From there, we calculated the distance from the centroid for all samples, and dropped samples that were outside of the 95^th^ percentile. This resulted in a total of 628 hiPSCs, comprising 359 female and 269 male hiPSCs.

As there were multiple clones per cell line for many individuals, we performed differential expression analysis after randomly selecting one unique cell line. To include all samples, we performed this random sampling 100 times. For each differential expression analysis, we filtered out low expression via filterByExpr, normalized for library sizes, before applying voom normalization (Law et al., 2014; Ritchie et al., 2015). Following voom normalization, we fitted a linear model (**Equation 1**) to examine sex. Differentially expressed genes were identified using the eBayes function (Smyth, 2004) from limma. For plots, we calculated the median differential expression statistics and voom expression across iterations.

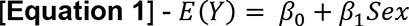

### Differential expression analyses and life span analysis of human postmortem brains

We downloaded FASTQ files and phenotype information for the caudate nucleus, dorsolateral prefrontal cortex (DLPFC) and hippocampus from the BrainSeq Consortium (Benjamin et al., 2020), selecting samples based on 1) diagnosis of neurotypical control and schizophrenia, 2) postnatal, 3) African and European ancestries, and 4) RiboZero RNA library preparation. This resulted in 1236 samples associated with 554 unique individuals (females, n = 174; males, n = 380). We aligned FASTQ files to GENCODE v26 with HISAT2 and generated sorted and indexed BAM files with SAMtools. Following alignment and sorting, we generated gene, repeat, and exon counts using featureCounts. For gene and repeat counts, we created a customized GTF file of genes and repeats by combining GENCODE v26 with repeat masker (UCSC Table Browser; hg38) repeat annotation after we annotated repeats by strand. With this GTF file, we then generated counts with featureCounts using six parameters: 1) paired end, 2) reversed stranded reads, 3) primary alignments only, 4) excluding chimeric reads, 5) allowing for multimapping reads and 6) one base as the minimum overlapping fraction in a read. For exon counts, we used GENCODE v26 with featureCounts for paired end, reversed stranded reads with four read filters: 1) uniquely mapped reads; 2) reads aligned in proper pairs; 3) excluding chimeric reads; and 4) primary alignments.

Similar to the hiPSC analysis, we assessed quality control based on read quality outliers determined via PCA and centroid analysis. We excluded samples that were outside of the 99^th^ percentile (caudate nucleus and hippocampus) and 95^th^ percentile (DLPFC), resulting in a total of 1203 samples: caudate nucleus (n=422), DLPFC (n=412), and hippocampus (n=369).

For life span differential expression analysis of human postmortem brains, we first selected neurotypical controls, resulting in a total of 768 samples (associated with 353 unique individuals): caudate nucleus (n=269), DLPFC (n=260), and hippocampus (n=239). After this sample selection, we then filtered out low expression with filterByExpr, and normalized for library sizes. For lifespan analysis, we used four age groups based on caudate developmental windows: A) 0-15 years, B) 16-30 years, C) 31-50 years, and D) 51+ years. For voom normalization, we fitted a linear model (**Equation 2**) for age group, adjusting for sex, ancestry (SNP PC 1-3 and self-reported), RNA integrity number (RIN), mitochondria mapping rate, gene assignment rate, genome mapping rate, and any hidden variances with surrogate variable analysis (Leek et al., 2012). We identified differentially expressed genes, repeats, and exons using eBayes. For plotting, we generated residualized data by regressing out covariates, and applying a z-score transformation from the voom normalized counts.

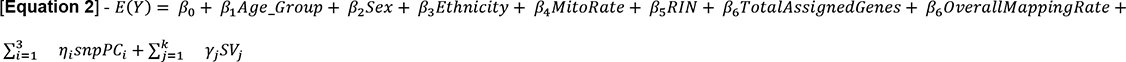

To examine sex differences for repetitive elements, we first selected adults (age > 17), resulting in a total of 1085 samples, associated with 479 unique individuals: caudate nucleus (n=394; 241CTL/153SZ; 120F/274M; 206AA/188EA), DLPFC (n=365; 213CTL/152SZ; 120F/245M; 202AA/163EA), and hippocampus (n=326; 196CTL/130SZ; 100F/226M; 178AA/148EA). After this sample selection, we then performed differential expression analysis by sex, for genes and repeats together, after filtering for low expression, normalizing for library size, and voom normalization. For voom normalization, we fit a linear model (**Equation 3**) for sex, adjusting for eight factors: diagnosis, age, ancestry (SNP PCs 1-3 and self-reported), RIN, mitochondria mapping rate, gene assignment rate, genome mapping rate, and hidden variance. For plotting, we generated residualized data using voom normalized counts by regressing out covariates and applying a z-score transformation.

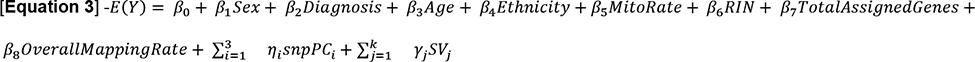

All transcriptomic analyses of *TAF1* expression at the gene and exon levels have been carried out referring to the canonical isoform *TAF1-204* (ENST00000423759.6). The sex differential expression analysis of *TAF1* gene expression in human postmortem brain tissues shown in Fig. 4F is a graphical representation of previously published data from the GTEX v8 collection (we referred to Supplementary Table S3 of (Oliva et al., 2020)).

### Dosage compensation analysis

For dosage compensation, we calculated relative X expression (RXE) as previously described (Jue et al., 2013) with slight modifications to replace FPKM (fragments per kilobase of transcript per million mapped reads) with TPM (transcripts per million). We generated TPM from gene counts using effective length calculated from mean insert size, which was determined using the Picard tool CollectInsertSizeMetrics (http://broadinstitute.github.io/picard/). Genes with an effective length less than or equal to 1 were excluded. To compute RXE, we filtered TPM genes with low expression log2 transformed prior to calculating the difference in the mean of autosome chromosomes compared to the X chromosome (**Equation 4**).

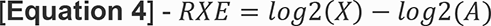

### SVA transcripts analysis

To detect SVA-derived transcripts in XDP female carrier iPSC lines, we aligned all reads to the XDP SVA sequence obtained from (Aneichyk et al., 2018) using HISAT2, and to calculate coverage at each base pair in SVA we used SAMtools.

### Graphics

We generated RXE scatter plots and boxplots using ggpubr (v0.4.0) in R.

### Code accessibility

All data are presented in the manuscript and the supplementary information. Code and jupyter notebooks are available through GitHub at https://github.com/apuapaquola.

## Results

### Identification of isogenic female carrier-derived iPSCs to model XDP

Genetically identical female iPSC lines that differ only in whether the mutant X chromosome or the wild-type X chromosome is expressed are a powerful tool for *in vitro* disease modeling. To model XDP by harnessing naturally occurring XCI, we characterized eight iPSC lines previously derived from three heterozygous female carrier donors (Aneichyk et al., 2018; Ito et al., 2016) (**Extended Data Fig.1-1a**). All eight XDP female carrier-derived iPSC lines carried the disease-specific haplotype on one of the X chromosomes, and we confirmed the 2.6 kb SVA retrotransposon insertion by long-range PCR amplification (Ito et al., 2016). The detection of two PCR products, of 0.6 kb and 3.2 kb, indicated that the samples were heterozygous for the S2control and XDP loci (**Extended Data Fig.1-1b**). Pluripotency analysis and BeadChIP genotyping array on all eight iPSC clones confirmed their primed pluripotent state, sample identity, and genomic integrity (**Extended Data Fig.1-2a-d**).

To evaluate the XCI status of the eight XDP carrier iPSC lines, we analyzed gene expression profiles and allele-specific expression from stranded total RNA-seq (median 30 M paired-end reads per clone) (Fig. 1a). To identify pairs of isogenic female iPSC lines with an identical genetic background, but expressing either the wild-type or XDP allele, we conducted an allele-specific expression analysis of expressed single nucleotide polymorphisms (SNPs) along the X chromosomes. For each set of clonal iPSC lines derived from the same female individual, we evaluated the percentage of X-linked monoallelically expressed SNPs that were shared or uniquely expressed between each clonal pair. When the same X chromosome is activated in two iPSC clones, most monoallelically expressed X-linked transcripts will be shared. In contrast, if different X chromosomes are activated, X-linked monoallelic transcripts will be expressed by different alleles in the two cell lines, and thus most SNPs will be uniquely expressed by each cell line.

**Figure 1.**
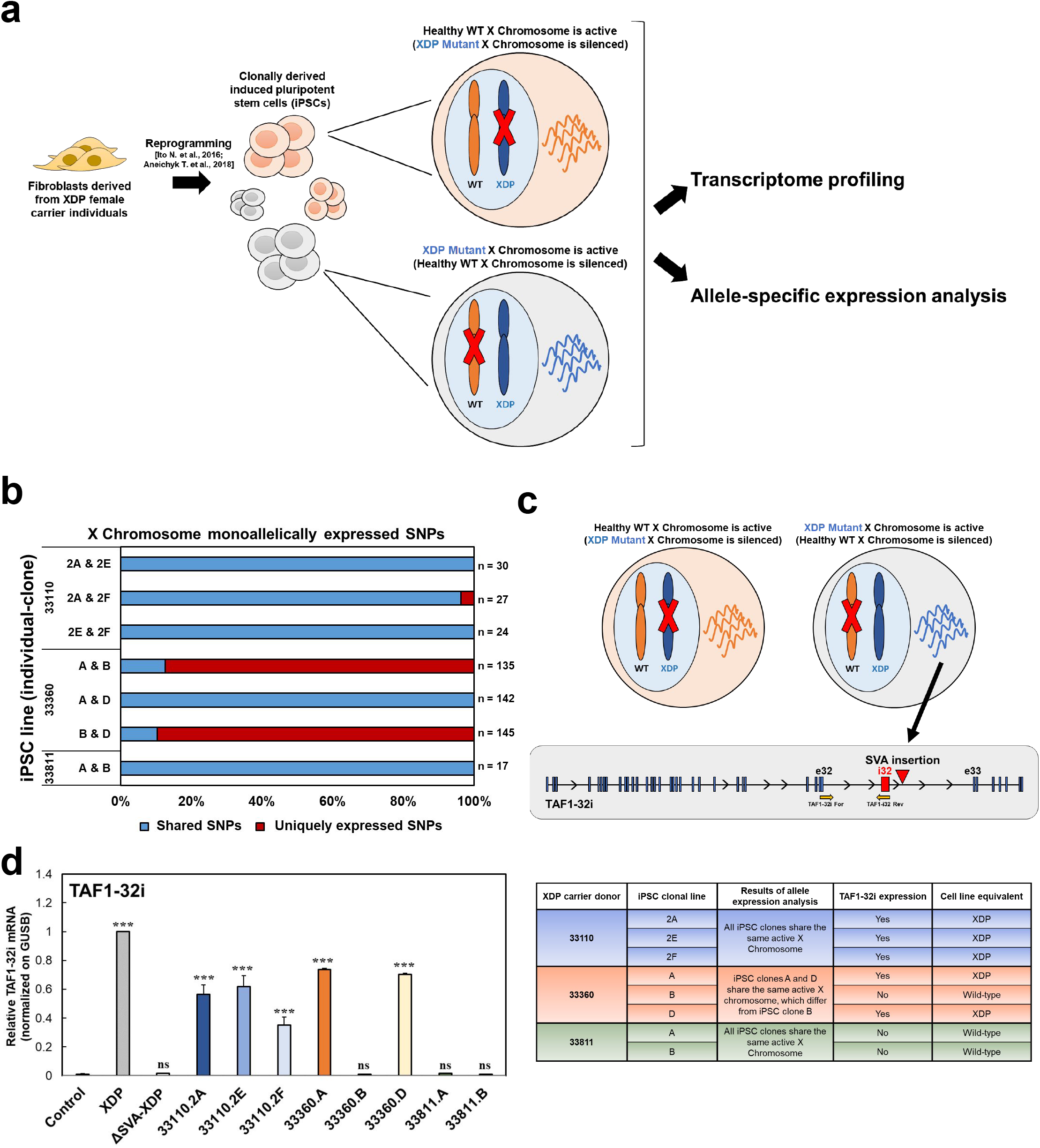
Characterization of XDP female carrier-derived iPSCs for X chromosome inactivation status of the XDP haplotype. **a.** Illustration of XDP female carrier-derived induced pluripotent stem cells (iPSCs). Due to clonal expansion of iPSCs and random XCI, female iPSC clones can inactivate two different X chromosomes. If the X chromosome carrying the healthy wild-type (WT) allele is active, the cell will express the canonical TAF1 transcript; in contrast, if the X chromosome carrying the mutant XDP allele is active, the cell will express the mutant TAF1 transcript, including the XDP haplotype. XDP female carrier-derived iPSCs were subjected to strand-specific RNA sequencing to perform gene expression analysis and allele-specific expression analysis. **b.** Bar plots of the fraction of allele-specific transcripts derived from the X-chromosomes, which are shared between the different iPSC clones. Expression from the same or different alleles was determined by counting the number of shared and unique homozygous single nucleotide polymorphisms (SNPs), respectively. The specific number of SNPs analyzed for each pair is shown to the right of the bar. **c.** Illustration of the mutant TAF1 splice variant characterized by retention of intron 32 (*TAF1-32i*) close to the SVA insertion. We used a designed custom TaqMan primer/probe to detect the presence of the exon 32/intron 32 splice site (Aneichyk et al., 2018). Yellow arrows show that forward and reverse primers bind to exon 32 and intron 32, respectively, so that the probe spans the splice junction. **d.** The graph shows the relative expression of *TAF1-32i* in XDP female carrier-derived iPSCs in comparison to iPSCs derived from control, XDP, and relatively isogenic SVA-deleted lines (ΔSVA-XDP). All values were normalized to the XDP sample. One-way ANOVA was performed on the mean and SEM from three independent experiments, and significance was determined as follows: *** p ≤ 0.001, ns=not significantly different. **e.** Table summarizing features of the eight XDP female carrier-derived iPSCs used in this study. *Figure contribution:* Ricardo S. Jacomini and Apua C.M. Paquola performed the allele-specific expression analysis; Laura D’Ignazio performed the TAF1-32i Taqman assay and analyzed the data.

We found that the three iPSC clones derived from the XDP female carrier 33360 were isogenic, with different active X chromosomes. Clones 33360.A and 33360.D shared the same active X chromosome, with all X-linked monoallelically expressed SNPs being shared. Clone 33360.B shared only 13% of monoallelically expressed SNPs with clone 33360.A and 10% with clone 33360.D, while 87% and 90% of X-linked monoallelically expressed SNPs were respectively uniquely expressed. This indicates 33360.B has a different active X chromosome compared to both clones 33360.A and 33360.D (Fig. 1b). Interestingly, clones 33360.A and 33360.D also exhibited high correlation with each other in the PCA analysis of normalized expression of X-linked genes from all XDP female carrier iPSC lines (**Extended Data Fig.1-3a**).

In contrast, the three iPSC clones derived from carrier 33110 shared the same active X chromosome, and >96% of X-linked monoallelically expressed SNPs. Similarly, all iPSC clones from carrier 33811 had the same active X chromosome, with all X-linked monoallelically expressed SNPs being shared (Fig. 1b).

To further validate that allele-specific expression indicated an active X chromosome from XCI, we analyzed the allele-specific expression of chromosome 7, an autosome equivalently sized to the X chromosome. In all iPSCs, we observed expression from the same allele for chromosome 7-specific genes, indicating monoallelic expression due to eQTLs or imprinting (**Extended Data Fig.1-3b**). Overall, the allele-specific expression analysis revealed that the iPSC clones derived from individual 33360 were isogenic XDP female carrier iPSC lines.

Next, we determined whether the active X chromosome in each XDP female carrier iPSC line represented the wild-type, or the XDP allele, by using total RNA-seq to quantify expression of the XDP-specific *TAF1-32i* transcript in the female carriers’ transcriptomic profiles (Fig. 1c). No significant differences were detected among the eight clones, likely due to the few transcripts mapped to the region, and our use of the total RNA-seq method (compared to capture sequencing used previously) (Aneichyk et al., 2018). Therefore, we used a *TAF1-32i*-specific TaqMan assay (Aneichyk et al., 2018) to measure the levels of this transcript in all female carrier iPSC lines. As expected, we detected *TAF1-32i* in XDP male iPSCs. We also detected *TAF1-32i* in 5 of the 8 female carrier-derived iPSC lines, with levels of expression ∼40-80% higher than those in the male control iPSCs (p < 0.001), and ∼35% lower than those in XDP patient-derived iPSCs. *TAF1-32i* was detected in clones 33360.A and 33360.D, but not in clone 33360. B, which was similar to the male control line (Fig. 1d). With this analysis, we defined the specific epigenetic identity of the isogenic XDP female carrier iPSC lines, with clone 33360. B being equivalent to wild-type cells, and clones 33360.A and 33360.D resembling XDP male cells (Fig. 1e).

### Characterization of X-Chromosome Inactivation in XDP female carrier-derived iPSCs

XCI is a dosage compensation mechanism that achieves equivalent X-linked gene expression between XY males and XX females. In the human female, two mechanisms ensure dosage compensation at different stages of development. In the pre-implantation embryo, X-linked genes are downregulated, termed dampened, along with coating by XIST of both chromosomes (Okamoto et al., 2011; Petropoulos et al., 2016; Sahakyan and Plath, 2016). Post-implantation, X chromosome inactivation occurs, such that one X chromosome is epigenetically silenced and lincRNA is coated by XIST on the inactive X chromosome. Female primed pluripotent stem cells generally harbor one active and one inactive X chromosome, but also demonstrate epigenetic variability in dosage compensation status, and some cultures demonstrate erosion of dosage compensation (Mekhoubad et al., 2012; Shen et al., 2008; Silva et al., 2008).

To assess epigenetic silencing of one X chromosome, we analyzed the total RNA-seq dataset for mono- and biallelic expression. Due to transcriptional silencing following X inactivation of one X chromosome, X-linked genes are expected to be transcribed mostly from one allele (monoallelic expression, with alternative allele frequency <0.1 or >0.9). In contrast, most autosomal genes are expressed biallelically from either maternal or paternal alleles, with alternative allele frequency ≈0.5. Therefore, when XCI properly occurs, the fraction of transcripts with monoallelic expression is expected to be greater for X-linked genes than for autosomal genes.

We quantified the number of monoallelically expressed transcripts from the X chromosome compared to all autosomal chromosomes, as the fraction of homozygously expressed SNPs for a chromosome, defined as the ratio of the number of expressed SNPs with an allele frequency <0.1 and >0.9 from a chromosome over the total number of expressed SNPs for the chromosome (**Extended Data Fig.1-3c**). For all eight iPSC lines, the fraction of X-linked monoallelically expressed transcripts was greater than that of autosomal genes, confirming that XCI occurs. The fraction of X-linked monoallelically expressed genes varied significantly among iPSC lines. Four out of eight iPSC lines (33110.2A, 33360.A, 33360.B, and 33360.D) demonstrated 4 to 6-fold greater monoallelically expressed X-linked transcripts compared to autosomal transcripts, indicating transcription from a single active X (XaXi status). The remaining four iPSC lines (33110.2E, 33110.2F, 33811.A, and 33811.B) exhibited only a 1.5 to 2-fold increase in monoallelically expressed X-linked transcripts compared to autosomal transcripts, indicating partial erosion of the inactive X chromosome (XaXe status) (**Extended Data Fig.1-3d**).

To investigate the expression dosage of X-linked genes on a chromosome-wide level, we compared the global transcriptional output of the X chromosome to that of autosomes. Briefly, we calculated the relative X expression (RXE), corresponding to chromosome-wide estimates of mean log-transformed TMP values of X-linked genes (X) compared to autosomal genes (A) (Duan et al., 2019). If the X and autosomal expression means are equal (RXE = 0), the average X-linked gene expression is comparable to the average expression from autosomal genes. If RXE > 0, the X chromosome is expressed at higher levels than the average autosome. RXE <0 and >-1 indicated higher autosomal gene expression than X (Fig. 2a).

**Figure 2.**
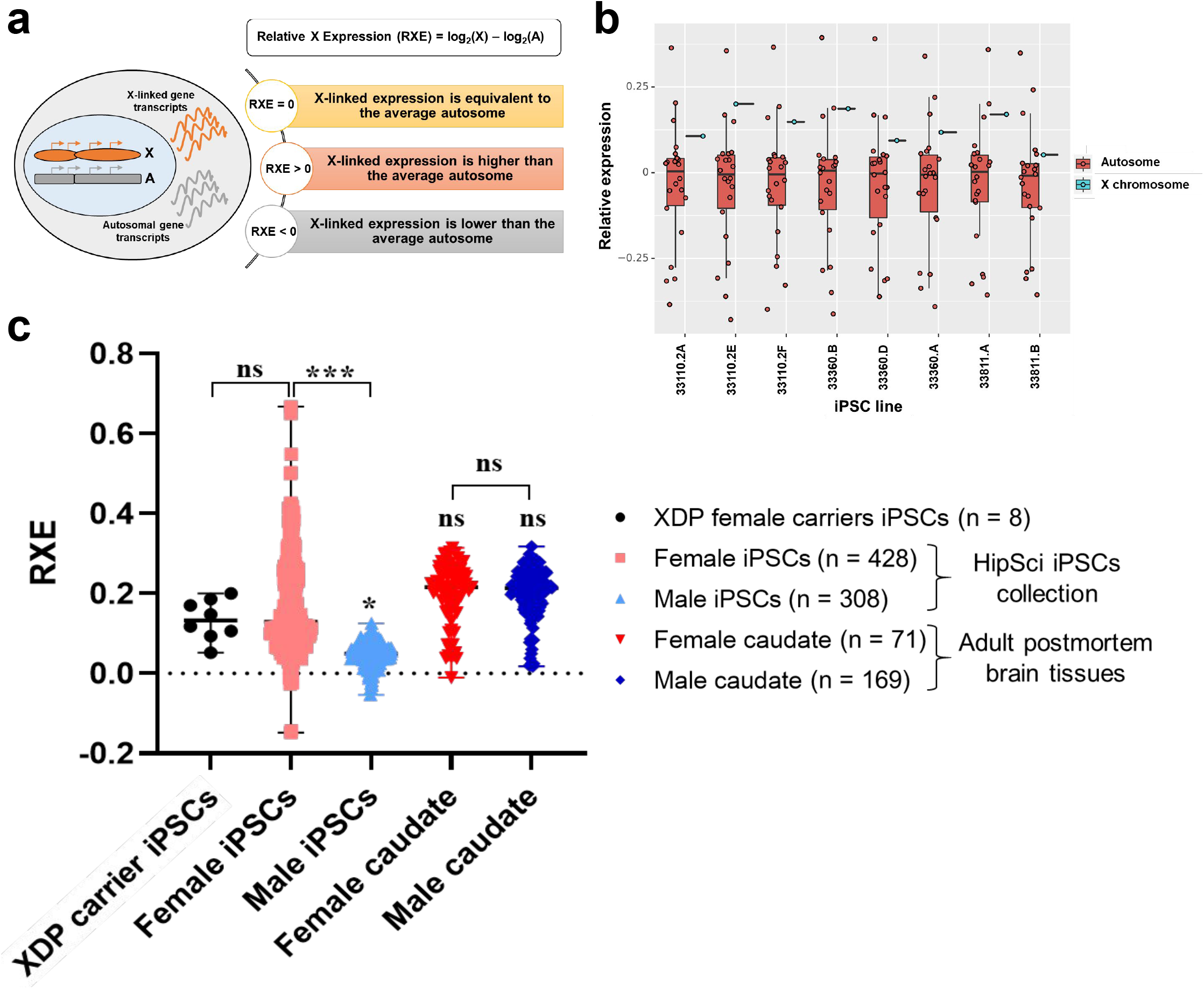
XDP female carrier-derived iPSCs have proper X chromosome dosage compensation. **a.** The relative expression of the X chromosome (RXE) is the difference between the log2-transformed mean TPM values of the X chromosome and all autosomes. If the expression of X and autosomes is equal, the RXE value will be 0, suggesting proper X dosage compensation. Positive RXE values indicate that the expression of X-linked genes is greater than that of autosomal genes; thus, dosage compensation is complete. Negative RXE values indicate that the expression of X-linked genes is lower than that of autosomes, and there is incomplete dosage compensation. **b.** Box plots showing log2-transformed relative expression of the X chromosome (RXE, blue) and all autosome chromosomes (red) for XDP female carrier-derived iPSCs. Red dots indicate the relative expression of each autosome pair (RGE) over all other chromosomes. **c.** RXE values of XDP female carrier-derived iPSC lines in comparison to female and male iPSC lines from public HipSci collection (Kilpinen et al., 2017), and adult female and male postmortem caudate nucleus samples from the BrainSeq Consortium (Benjamin et al., 2020; Collado-Torres et al., 2019). One-way ANOVA was performed on the mean and SEM calculated for each sample, and significance was determined as follows: * p ≤ 0.05, *** p ≤ 0.001, ns=not significant. *Figure contribution:* Apua C.M Paquola and Kynon J.M. Benjamin performed the dosage compensation analysis.

In all eight XDP female carrier-derived iPSCs, RXE values ranged from 0.05 to 0.20, suggesting similar levels of X-linked gene dosage in all lines (Fig. 2b). No statistically significant difference (p = 0.8399) was observed when comparing the median RXE of the eight XDP female carrier iPSCs (median RXE = 0.1344) with the median RXE of the large publicly available HipSci collection of wild-type female iPSC lines (median RXE = 0.1662; n = 438 (Kilpinen et al., 2017)). Aberrant eroded iPSC lines in the HipSci collection demonstrated an RXE value of up to 0.6, a value much higher than that of the XDP iPSCs. The RXE analysis demonstrated that the XDP carrier iPSC lines have X chromosome dosage compensation levels similar to those of most wild-type female iPSC lines. These HipSci lines maintain an inactive X (Bar et al., 2019). The median RXE of male iPSCs (median RXE = 0.0482; n = 308, (Kilpinen et al., 2017)) was significantly lower than that of female iPSCs (p = 0.0428 compared to XDP carrier iPSCs; p < 0.001 compared to female HipSci iPSCs), due to genes escaping XCI.

To measure dosage compensation in a postmortem brain tissue-specific context, we calculated the RXE for caudate nucleus samples (Benjamin et al., 2020; Collado-Torres et al., 2019), and found that median RXE values in both female (median RXE = 0.1985; n = 71) and male (median RXE = 0.2048; n = 169) caudate samples were slightly higher than those in wild-type female iPSCsfrom HipSci collection, without reaching statistical significance. No statistically significant difference was detected between female and male caudate samples (p = 0.9862), suggesting full dosage compensation (Fig. 2c). In summary, the isogenic lines 33360.A, 33360.B and 33360.D demonstrated proper XCI with monoallelic expression and dosage compensated X-linked gene expression.

### Isogenic XDP female carrier iPSC lines display variable XIST expression

Two long noncoding RNAs (lncRNAs), *XIST* and *XACT,* regulate human XCI. *XIST* (X-inactive coating transcript) coats the inactive X chromosome *in cis,* triggering gene silencing through epigenetic changes (Cantone and Fisher, 2017). *XACT* (X-active coating transcript) coats the active X chromosome in early preimplantation development (Vallot et al., 2013).

In the total RNA-seq data from the XDP female carrier-derived iPSCs, the X-linked transcripts *XACT*, *ATRX* and *HUWE1* were expressed at similar levels among all iPSC lines (Fig. 3a). *TSIX*, the antisense transcript that negatively regulates *XIST (Stavropoulos et al., 2001)* in mice (but with unknown function in humans), was detected at extremely low levels in all iPSCs. Consistent with previous reports, *XIST* expression varied significantly, and even iPSC clones derived from the same XDP carrier individual exhibited statistically significant differences in *XIST* expression (Fig. 3a-b). Two out of eight iPSC lines, 33110.2E and 33360.B, exhibited minimal *XIST* mRNA expression that was comparable to that of a male iPSC line. The iPSC clones 33360.A and 33360.D, the mutant-equivalent cell lines of the isogenic set, showed *XIST* levels ∼45-60% higher than the wild-type equivalent iPSC clone 33360.B. Our previous analyses demonstrated that monoallelic expression and dosage compensation occurred in all XDP female carrier-derived iPSC lines, despite low levels of *XIST* (Fig. 2 and **Extended Data Fig.1-3**). The *XIST-*primed iPSC state is thought to be an abnormal state that can proceed with erosion of X chromosome silencing; however, we demonstrate that global silencing of X was preserved, despite low levels of *XIST*.

**Figure 3.**
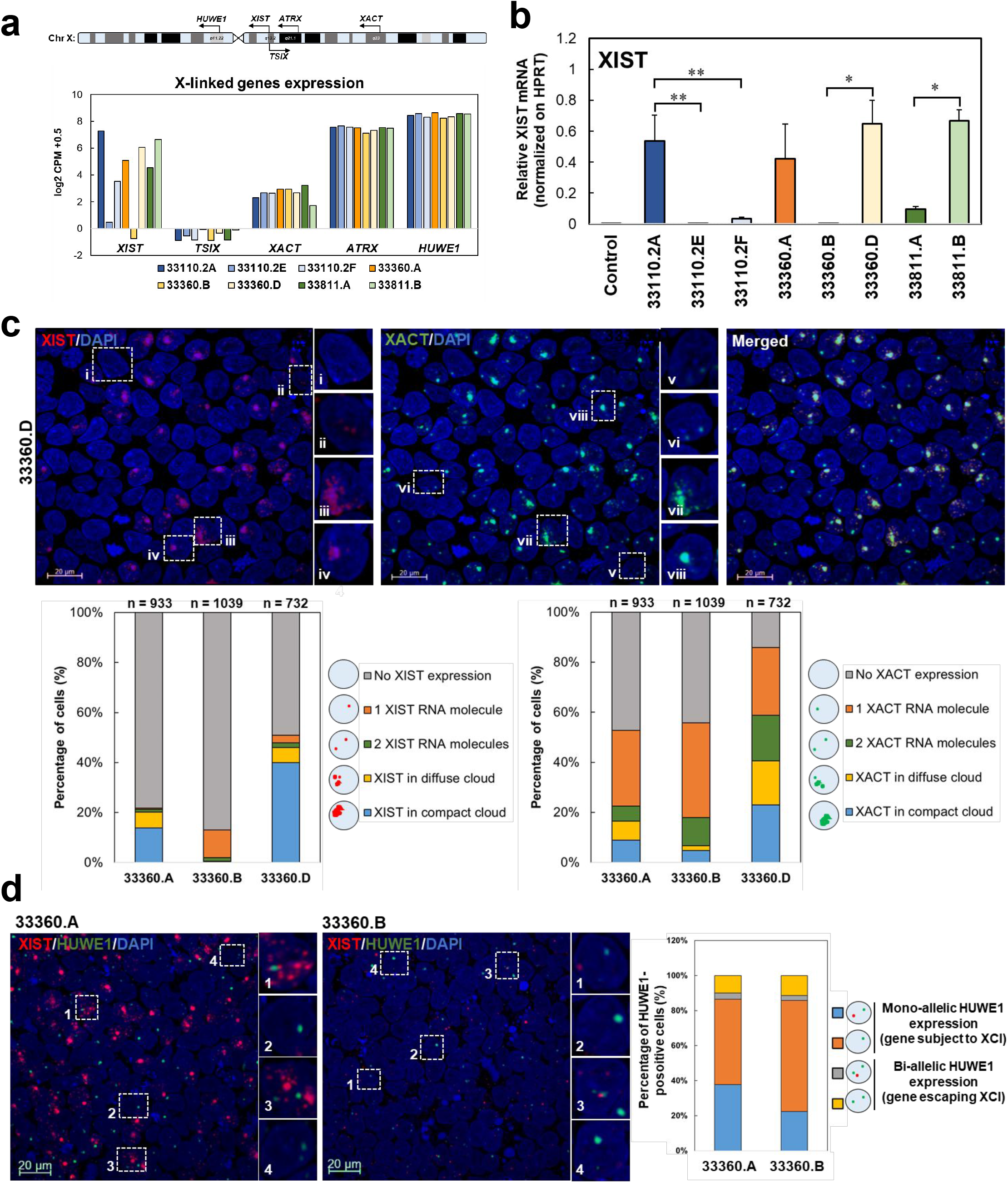
Dosage compensation in female iPSCs inversely correlates with *XIST* expression. **a.** Simplified map of the *XIST*, *TSIX*, *HUWE1*, *ATRX*, and *XACT* loci on the X chromosome, and expression of X-linked gene transcripts in XDP carrier-derived iPSCs. Out of these genes, *XIST* shows higher variability among clones. **b**. RT-qPCR analysis of *XIST* expression in XDP female carrier-derived iPSCs compared to a control male iPSC line. Primers flanking exons 1 and 3 of XIST have been used. One-way ANOVA was performed on the mean and SEM from three independent experiments, and significance was determined as follows: * p ≤ 0.05; ** p ≤ 0.01. **c.** Representative images from iPSC clone 33360.D showing localization of RNA scope-specific probes for *XIST* (red) and *XACT* (green) transcripts. DAPI was used to stain nuclei. Scale bar is 20 µm. Based on the expression of those transcripts, different categories of nuclear localization were identified: i/v) detection of one RNA molecule of *XIST/XACT*; ii/vi) detection of two RNA molecules of *XIST/XACT*; iii/vii) detection of several dispersed molecules of *XIST/XACT* transcripts forming a diffuse cloud; iv/viii) detection of several molecules of XIST/XACT transcripts organized in a compact cloud. Bar charts report quantification of the four categories in the isogenic set of XDP female carrier-derived iPSCs. For each clone, > 700 cells have been analyzed. The percentage of cells not expressing XIST or XACT transcripts is also reported. **d.** Representative images from iPSC clone 33360.A and 33360.B showing localization of RNA scope-specific probes for XIST (red) and HUWE1 (green) transcripts. DAPI was used to stain nuclei. Scale bar is 20 µm. The HUWE1 gene is usually subject to XCI; thus, four different cellular conditions were identified: 1) monoallelic expression of HUWE1 accompanied by XIST expression (in any form, i.e., single RNA molecules, multiple RNA molecules, diffuse clouds, compact cloud); 2) monoallelic expression of HUWE1 in the absence of XIST expression; 3) biallelic expression of HUWE1 accompanied by XIST expression (in any form, i.e., single RNA molecule, multiple RNA molecules, diffuse cloud, compact cloud); 4) biallelic expression of HUWE1 in the absence of XIST expression. Bar graphs report the percentage of HUWE1-positive cells showing each of the four cellular conditions in the isogenic iPSC pair 33360.A and 33360.B. For each clone, > 600 cells have been analyzed. *Figure contribution:* Kynon J.M. Benjamin analyzed the RNA-seq data; Laura D’Ignazio and Bareera Qamar performed and analyzed both the XIST qPCR analysis and the RNA scope.

While the transcriptomic data demonstrated that X-linked dosage compensation takes place at the population level of cells, we further assessed XCI at both single-cell and single-molecule resolution by using RNAscope *in situ* hybridization. Through canonical RNA FISH, *XIST* is usually visualized as small puncta or a single large cloud accumulating on the inactive X chromosome (Erwin and Lee, 2010). RNAscope technology allows for multiplex detection of target RNAs at a single-molecule resolution, with increased specificity and sensitivity compared to conventional multigene RNA-FISH that is detected using BAC probes (F. Wang et al., 2012). Here, we identified four different patterns of *XIST* nuclear localization: i) expression of one *XIST* RNA molecule when only a single *XIST*-specific fluorescent punctum was detectable; ii) expression of two *XIST* RNA molecules when two distinct *XIST*-specific fluorescent puncta were observed; iii) expression of multiple *XIST* RNA molecules forming a diffuse cloud when more than three dispersed *XIST*-specific fluorescent puncta were detected; and iv) expression of *XIST* as one condensed cloud (Fig. 3c**, Extended Data Fig.3-1a**). Due to the high resolution of RNAscope, allowing single-molecule visualization of specific RNAs of interest, there are often additional dispersed *XIST* molecules in cells with one condensed cloud. For our quantitative analysis, these cells were included in the fourth category, one condensed cloud. *XIST* expression and localization varied among the three isogenic iPSC clones, being detected in 22% of 33360.A iPSCs, 13% of 33360.B iPSCs, and 51% of 33360.D iPSCs. In iPSC clone 33360.B, 85% of cells exhibiting *XIST* expression expressed only one single *XIST* RNA molecule. In contrast, in iPSC clone 33360.A, 29% and 63% of *XIST*+ cells exhibited *XIST* expression in diffuse clouds or compact clouds, respectively. Among cells showing *XIST*expression in iPSC clone 33360.D, 12% exhibited *XIST* expression in diffuse clouds, and 72% displayed *XIST* in compact clouds (Fig. 3c). These variations might be correlated with differences in the total *XIST* expression observed at the transcriptomic level.

Interestingly, we detected also four patterns of *XACT* nuclear localization: (*i*) one single RNA molecule, (*ii*) two distinct RNA molecules, (*iii*) multiple dispersed RNA molecules organized in a diffuse cloud, and (*iv*) one large compact cloud was also detected using an *XACT*-specific fluorescent probe (Fig. 3c, **Extended Data Fig.3-1a**). A small fraction of cells contained one condensed XACT cloud along with other dispersed smaller *XACT* molecules or one less intense *XACT* molecule. These cells were counted in the category of one condensed cloud. *XACT* coated 53% of 33360.A iPSCs, 56% of 33360.B iPSCs, and ∼85% of 33360.D iPSCs. Nuclear colocalization of *XIST* and *XACT* was observed in all cells of iPSC clones 33360.A and 33360.D (Fig. 3c, **Extended Data Fig.3-1b**). In contrast, in most cells from clone 33360.B, *XIST* and *XACT* had distinct nuclear localization (**Extended Data Fig.3-1b**). Here, we report that the coaccumulation of *XIST* and *XACT* on the same X chromosome occurs not only in naive hESCs (Vallot et al., 2017), but also in human primed iPSC cultures.

To further evaluate X chromosome silencing at a single-cell resolution, we evaluated the single-molecule expression and localization of *HUWE1,* an X-linked gene subject to XCI, relative to *XIST* (Patel et al., 2017). If the X chromosome is silenced, *HUWE1* will be expressed from one X chromosome (monoallelic expression). Alternatively, if XCI mechanisms are impaired, *HUWE1* will be expressed by both X chromosomes (biallelic expression). For iPSC clones 33360.A and 33360.B, >80% of iPSCs demonstrated monoallelic expression of *HUWE1*, suggesting proper XCI (Fig. 3d); only 13% of 33360.A iPSCs and 10% of 33360.B iPSCs displayed biallelic expression of *HUWE1*. Approximately 49% of 33360.A iPSCs and 60% of 33360.B iPSCs monoallelically expressed *HUWE1* in the absence of *XIST* expression (Fig. 3d).

In summary, our results highlight the significant variations in XIST and XACT localization in human primed iPSC lines, as has been described previously. These findings demonstrate that X chromosome silencing and X-linked gene dosage compensation occur in isogenic XDP female carrier iPSC lines, independent of persistent *XIST*coating.

### Sexually dimorphic expression of TAF1 in pluripotent stem cells

Canonical *TAF1* expression was decreased in various XDP samples, including fibroblast, RNA from blood and NSCs, compared to control samples (Al Ali et al., 2021; Aneichyk et al., 2018; Domingo et al., 2016; Ito et al., 2016). To quantify the 3’ region of canonical *TAF1* expression in XDP female carrier-derived iPSCs, we performed RT-qPCR with primers targeting *TAF1* exons 32 and 33 (Rakovic et al., 2018).

Unexpectedly, all eight XDP female carrier iPSC lines displayed similar levels of expression of the canonical full-length *TAF1* transcript (Fig. 4a), regardless of their expression of the mutant isoform *TAF1-32i* (Fig. 1c). Next, we compared *TAF1* expression in XDP female carrier lines to that in male control- and XDP-derived iPSCs, and XDP patient-derived iPSCs where the SVA was excised, ΔSVA-XDP (Aneichyk et al., 2018) (Fig. 4b). As expected, *TAF1* mRNA levels in XDP patient-derived iPSCs were approximately 50% lower than in control cells (p < 0.001), and the expression normalized with SVA removal. Seven of the eight female carrier iPSC lines displayed a significant reduction in *TAF1* expression, compared to a control male iPSC line, with levels comparable to those of XDP iPSCs (Fig. 4b).

**Figure 4.**
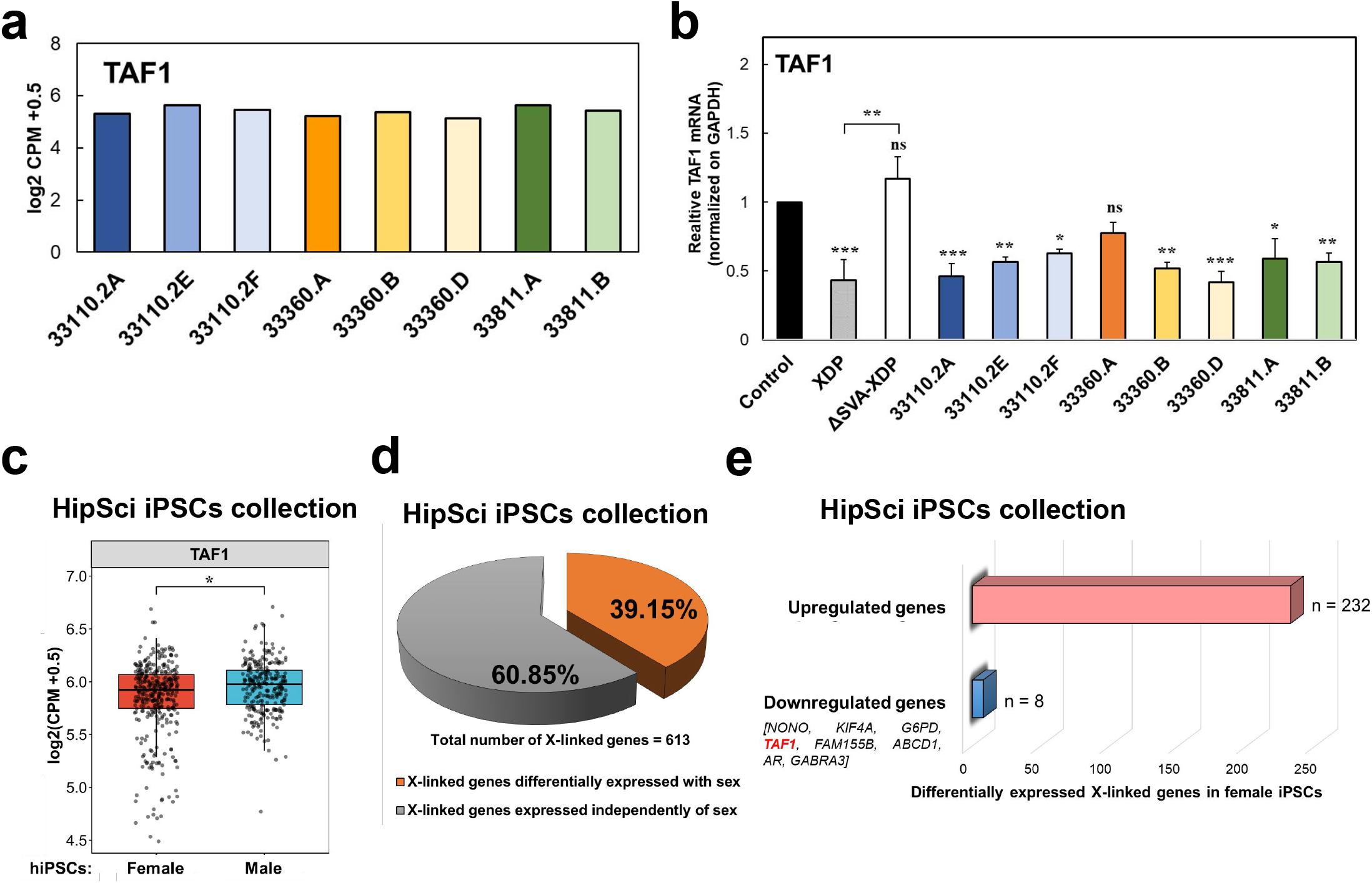
Sex-dependent expression of TAF1 in female iPSCs. **a.** Normalized expression of the TAF1 gene in XDP carrier-derived iPSCs. **b.** RT-qPCR analysis of TAF1 expression in XDP female carrier-derived iPSCs compared to iPSCs derived from control, XDP, and a relatively isogenic SVA-deleted line (ΔSVA-XDP). Primers flanking exons 32 and 33 of TAF1 were used in this analysis. One-way ANOVA was performed on the mean and SEM from three independent experiments, and significance was determined as follows: * p ≤ 0.05; ** p ≤ 0.01; *** p ≤ 0.001, ns = non-significant. **c.** Box plots comparing normalized TAF1 transcript expression levels in female (red) and male (blue) iPSCs from public HipSci collection **(Kilpinen et al., 2017)**. * indicates a p value ≤ 0.05. **d**. Percentage of X-linked genes in female iPSCs **(Kilpinen et al., 2017)** differentially expressed with sex (orange) and expressed independently of sex (gray). **e.** Number of sex-differentially expressed X-linked genes that are upregulated or downregulated in female iPSCs (Kilpinen et al., 2017). TAF1 is among the eight genes underexpressed in female iPSCs compared to male iPSCs. *Figure contribution:* Laura D’Ignazio performed the TAF1 qPCR analysis. Laura D’Ignazio and Kynon J.M. benjamin analyzed the RNA-seq data from XDP female carrier-derived iPSCs and HipSci collection.

Decreased *TAF1* expression in carrier iPSCs could be caused by either an XDP female carrier iPSC molecular phenotype, or a sexually dimorphic expression in all female iPSCs. To distinguish between these two possibilities, we analyzed the publicly available iPSC transcriptome dataset from 301 individuals (Kilpinen et al., 2017). *TAF1* was significantly decreased in wild-type female iPSCs compared to male cells (p = 0.0422) (Fig. 4c), indicating a sexually dimorphic expression of *TAF1*. X-linked genes are rarely decreased in females. A total of 240 X-linked genes were differentially expressed between sexes in human iPSCs (Fig. 4d). Eight genes, including TAF1, were downregulated in female iPSCs, while 232 of these differentially expressed genes were upregulated (logFC = 0.10201; p = 0.0422) (Fig. 4e). In summary, *TAF1* was decreased in control female iPSC lines, and decreased female expression was uncommon for X-linked genes.

### Striatal regions have equal levels of *TAF1* expression in males and females

To evaluate whether *TAF1* expression was decreased in the brain of adult females compared to males, we further investigated the recently reported sex expression differences in adult brains (Oliva et al., 2020). In the GTEx analysis, *TAF1* expression was significantly decreased (with LFSR<0.05) in 26 female tissues compared to males, including 8 brain regions, such as amygdala, anterior cingulate cortex, cortex, frontal cortex, hippocampus, hypothalamus, spinal cord, and substantia nigra (Fig. 5a). Importantly, no significant difference in *TAF1* expression between females and males was observed in the caudate nucleus, putamen or nucleus accumbens.

**Figure 5.**
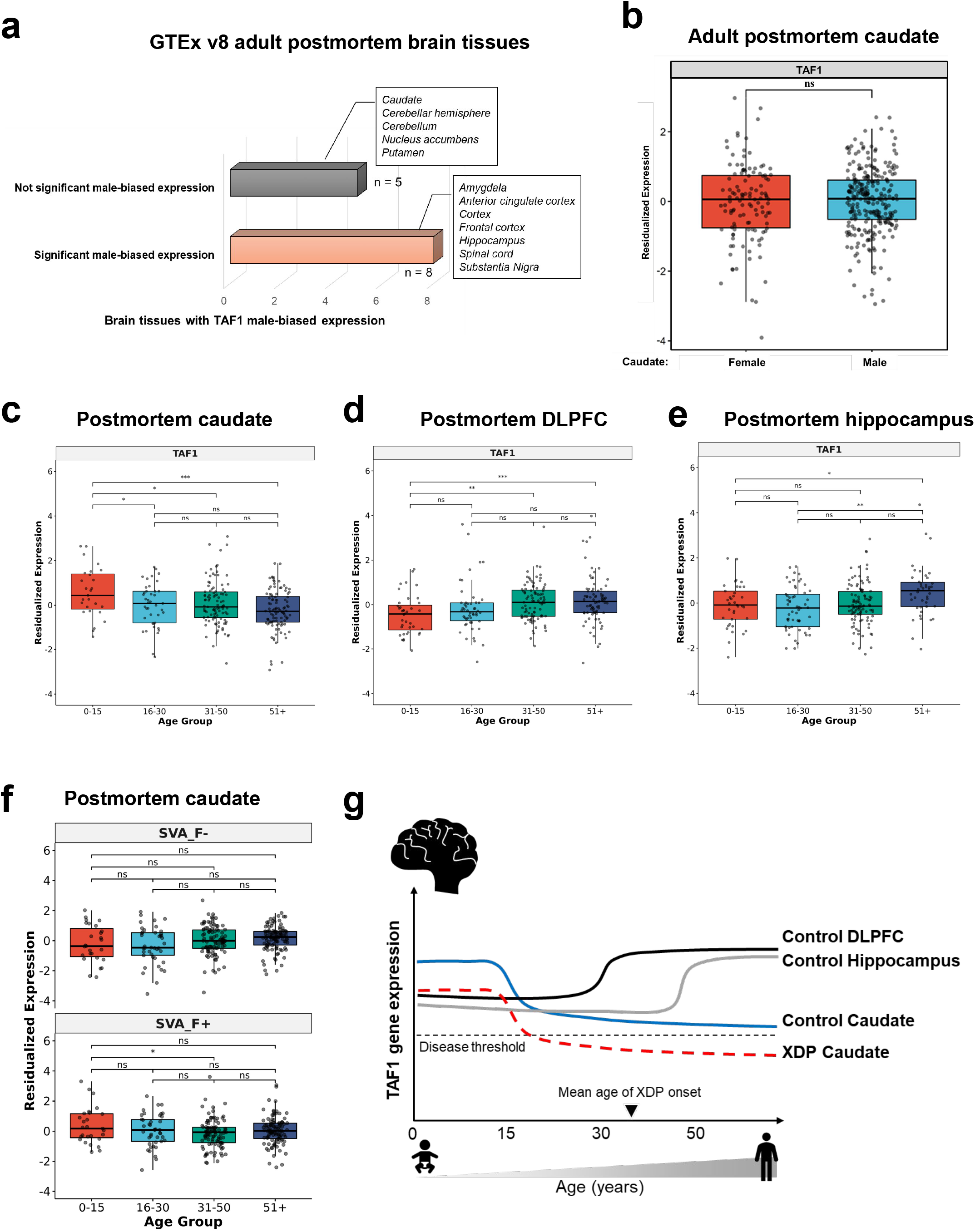
TAF1 and SVA-F expression over the lifespan in the postmortem brain. **a.** Number of postmortem brain tissues included in the GTEX v8 collection (Oliva et al., 2020) in which TAF1 expression is significantly or not significantly male biased. **b.** Box plots comparing residualized gene expression of TAF1 in the caudate nucleus (n = 394) of female (red, n = 120) and male (blue, n = 274) individuals from the BrainSeq Consortium (Benjamin et al., 2020; Collado-Torres et al., 2019). ns = not significant. **c-f.** Box plots comparing residualized gene expression of individuals from the BrainSeq Consortium (Benjamin et al., 2020; Collado-Torres et al., 2019) **(c)** TAF1 in the caudate nucleus, **(d)** TAF1 in the dorsolateral prefrontal cortex (DLPFC), **(e)** TAF1 in the hippocampus, **(f)** for SVA-F in the caudate nucleus. Samples were divided into four groups based on the age of the individuals: 0-15 years; 16-30 years; 31-51 years; and older than 51 years. ns = not significant; * FDR ≤ 0.05, **FDR ≤ 0.01, *** FDR ≤ 0.001. **(g)** Diagram depicting our proposed model to explain the differential degeneration in the neostriatum of individuals with XDP. The decrease in *TAF1* expression after 15 years of age may synergize with a XDP-specific partial loss of *TAF1* function in the caudate nucleus, a brain region that seems to be selectively sensitive to *TAF1* levels. *Figure contribution:* Laura D’Ignazio analyzed the publicly available GTEx dataset; Ria Arora performed the sex differential expression of TAF1; Kynon J.M. Benjamin performed the age-differential expression of TAF1 across lifespan; Taylor Evans performed the SVA-F differential expression analysis across lifespan.

We further evaluated *TAF1* expression in neurotypical controls included in the BrainSeq Consortium dataset, and confirmed that *TAF1* is not sexually dimorphic in the caudate nucleus (Fig. 5b). In the BrainSeq Consortium dataset, *TAF1* was not significantly sexually differentially expressed in the hippocampus or dorsolateral prefrontal cortex, (a finding which contrasted with the decreased female expression in the GTEX samples, described in the preceding paragraph, possibly due to sample variation in age or analysis pipelines) (**Extended Data Fig.5-1a**). As a positive control, a significant increase in *XIST* expression was consistently observed between female and male tissue samples (p < 0.001) (**Extended Data Fig.5-1b**).

In summary, *TAF1* was decreased in most wild-type female cells and tissues compared to males, including wild-type iPSC cells, mutant carrier XDP iPSC cells and female postmortem brains. However, *TAF1* was expressed equally in males and females in caudate nucleus, putamen, and nucleus accumbens, indicating region-specific regulation of *TAF1*.

### *TAF1* is decreased in adult postmortem caudate compared to young individuals

Next, we investigated *TAF1* expression in postmortem brain tissue over the lifespan, using recent lifespan neurogenomic datasets from the BrainSeq Consortium (Collado-Torres et al., 2019; Fromer et al., 2016; GTEx Consortium et al., 2017). By analyzing the total RNA-seq dataset of caudate nuclei from neurotypical individuals aged 0-90 years (Benjamin et al., 2020; Collado-Torres et al., 2019), we found that *TAF1* gene expression significantly decreased after 15 years of age (Fig. 5c). Individuals aged 0-15 years had the highest level of *TAF1* expression compared to all other age groups (16-30 (FDR<0.05), 31-50 (FDR<0.05), and 51+ years (FDR<0.0005)) (**Extended Data Table 5-1**). Notably, increased *TAF1* expression occurred throughout the transcript. When examining *TAF1* expression at the exon level, 25 out of 38 exons located throughout the entire transcript were significantly increased in young individuals. The remaining 13 exons were excluded from this analysis due to their low expression (below the threshold for accurate quantification) (**Extended Data Table 5-2**).

*TAF1* expression demonstrated an opposite age association in the DLPFC and hippocampus. In the DLPFC, there was higher expression in 31- to 50-year-olds and 51+ compared to 0- to 15-year-olds (Fig. 5d). In the hippocampus, TAF1 expression was higher in the 51+ age group compared to younger groups (Fig. 5e). Considering the reported onset of XDP at approximately age 35, the specific decrease in *TAF1* expression after 15 years of life in the caudate nucleus precedes the age of onset of symptoms (Fig. 5g). Taken together, these findings support the existence of age-, tissue- and sex-specific fine regulatory mechanisms controlling *TAF1* expression.

### Evaluating SVA-F expression

In addition to variation in *TAF1* expression, regulation of SVA expression in the brain may also influence the XDP phenotype. While neurotypical brains do not have the patient specific SVA-F insertion in TAF1 intron 32, other SVA-F copies with identical sequences are present in all humans and these sequences are regulated by cellular host factors in trans. Therefore, we evaluated SVA-F expression in general in postmortem brain samples from BrainSeq, which employs a stranded total RNA-seq protocol. Because SVA-F elements are repetitive and have identical sequences, we quantified all transcripts mapping to any SVA-F location in the genome, the human-specific subfamily involved in XDP (Hancks and Kazazian, 2010).

The expression of the SVA-F retrotransposon was not sexually dimorphic in the caudate nucleus of healthy individuals from the BrainSeq dataset (**Extended Data Fig.5-2a**). In general, SVA-F plus strand and minus strand expression showed slight or no significant difference between most age groups for the caudate nucleus, DLPFC and hippocampus (Fig. 5f**, Extended Data Fig.5-2b-c)**. No significant differences were seen for any comparison of SVA-F minus strand expression. In the caudate nucleus and DLPFC, SVA-F plus strand expression was slightly but significantly decreased in 31- to 50-years-old individuals, compared to 0- to 15-years-old samples (FDR <0.01); In DLPFC, decreased SVA-F plus strand expression was also significantly decreased in 51+ (FDR<0.01) samples, compared to 0- to 15-year-olds group. In summary, we found abundant expression of SVA-F in neurotypical brains, slightly decreased SVA-F expression in caudate nucleus and DLPFC after 30 years of life and no differential expression of SVA-F between sexes.

Transcripts from the XDP-specific SVA and the hexanucleotide repeat could drive neurodegeneration in XDP. To evaluate the potential for disease-associated SVA-derived transcripts, we analyzed SVA expression in XDP female carrier iPSCs. We identified abundant SVA-derived transcripts (**Extended Data Fig.5-2d**). However, due to the multicopy nature of SVA-F, it was unclear whether these transcripts arose from XDP-specific SVA or another SVA-F. Most transcripts aligned to the SINE-R and Alu-like regions in both sense and antisense orientation, and aligned equally well to the XDP-specific SVA-F and approximately 4-10 other SVA-F regions in the genome. In addition, transcripts were not significantly differentially expressed between wild-type- or mutant-equivalent XDP carrier iPSCs (**Extended Data Fig.5-2d**).

To detect XDP-specific SVA transcripts, we created a modified reference genome by introducing the XDP-specific SVA sequence at its corresponding position in chromosome X in hg38. We aligned the RNA-Seq reads to this modified reference genome, and assessed sequencing reads that aligned to the XDP-specific SVA. We found weak evidence for XDP-specific SVA-derived transcripts. In one mutant iPSC sample, one sequencing read spanned the Alu-like region to the hexamer repeat, and preferentially aligned to the XDP-specific SVA, due to a 1 bp difference unique to the XDP SVA compared to all other SVA-Fs. No transcripts or junctions were detected that spliced into the SVA from the TAF1 region. We attribute the weak evidence for XDP-specific SVA-derived transcripts, to the scarcity of transcripts and the redundancy of the sequence. In summary, we found abundant expression of SVA-F in iPSC lines regardless of XDP status, and weak evidence of XDP-specific SVA transcripts in iPS cells.

## Discussion

Our results demonstrate significant sex-, age- and tissue-specific differences in XDP-associated transcript expression in female iPSCs and in human postmortem brain. *TAF1* was moderately decreased in most female cells and tissues compared to males, including wild-type iPSCs, carrier XDP iPSCs, and in several regions of the female postmortem brain; *TAF1* was expressed equally in the male and female striatum. *TAF1* expression uniquely decreased after 15 years of age in the caudate nucleus but not in the hippocampus or DLPFC. SVA-F plus strand expression slightly decreased in caudate nucleus and DLPFC after 30 years of life in neurotypical brain, but we find weak evidence for XDP-specific SVA derived transcripts in female iPSC cells. These findings indicate that regional-, age- and sex-specific mechanisms regulate *TAF1* and SVA-F expression, further highlighting the importance of disease-relevant cellular models and postmortem tissue analysis.

### Analysis of XCI status is crucial for in vitro disease modeling

As a valuable cell resource for disease modeling, we extensively characterized eight iPSC lines from XDP female carrier individuals, and identified isogenic lines where one clonal iPSC line expressed the wild-type X, and the two other clonal iPSC lines expressed the XDP haplotype. In our study, we provided strong evidence that monoallelic expression and dosage compensation occur in all XDP female carrier-derived iPSCs here examined, independently of XIST RNA levels or XIST/XACT nuclear location patterns. This finding is consistent with previous analyses conducted on large cohorts of female iPSC lines and demonstrating monoallelic X-linked expression and dosage compensation for the majority of X-linked genes (Shen et al., 2008; Silva et al., 2008; Mekhoubad et al., 2012; Vallot et al., 2013; Cantone and Fisher, 2017; Bar et al., 2019; Patrat et al., 2020).

The variability of XCI status in primed iPSCs remains a significant challenge for female stem cell models. Dosage compensation state for primed pluripotency comprises: X chromosome inactivation with XIST coating a single silenced X, heterochromatinization of the silent X, and monoallelic expression for most X linked genes (Mekhoubad et al., 2012; Shen et al., 2008; Silva et al., 2008). The XCI state often erodes in primed pluripotent human ES cells, such that portions of the previously inactive X chromosome become reactivated and are not dosage compensated (Mekhoubad et al., 2012; Shen et al., 2008; Silva et al., 2008). The naive state is a pluripotent state that captures an earlier developmental state and can reset the XCI state of primed human stem cells (Sahakyan et al., 2017). However, the naive state is characterized by increased genomic instability and altered imprinting (Theunissen et al., 2016), and also require additional culture optimization to provide more suitable cells for disease modeling.

Differentiating the set of isogenic XDP female carrier-derived iPSC into specific neural lineages will help elucidate the molecular mechanism causing XDP, as these models may capture early stages of disease no longer present in end-state postmortem analysis. It has consistently been demonstrated that the differentiation of iPSC lines maintains the dosage compensation status during differentiation (Mekhoubad et al., 2012; Silva et al., 2008). We anticipate that the XDP female carrier iPSC lines described in this study should maintain XCI during neural differentiation. Importantly, we demonstrate absence of XIST expression in the majority of female carrier primed iPSCs, which indicates that the inactive X chromosome may further erode in some cultures. Therefore, careful analysis of the XCI status of future iPSC-derived neuronal models will be essential.

### XDP in female carriers and isogenic carrier-derived iPSCs

In this study, we characterized eight iPSC lines obtained from three XDP heterozygous female carriers (Ito et al., 2016; Aneichyk et al., 2018). Our analyses demonstrated that clonal carrier-derived iPSC lines containing the silent wild-type, and the active mutant allele, retain molecular features of XDP mutant lines, such as the expression of the XDP-associated aberrant transcript TAF1-32i. In contrast, female lines containing the active wild-type allele and silenced mutant allele had no detectable levels of TAF1-32i, a situation similar to control male iPSC lines and lines with a CRISPR-deleted SVA. A recent study demonstrated that TAF1-32i expression was elevated in XDP patients and female carriers compared to control samples but TAF1-32i expression levels did not distinguish carriers vs affected XDP patients (Al Ali et al., 2021). Our data suggest that the XDP haplotype is fully silenced by XCI and that TAF1-32i expression is a biomarker for XDP haplotype expression in female pluripotent stem cells. Future investigations will be needed to determine if TAF1-32i expression is a biomarker for XDP haplotype expression in carrier samples and differentiated cells.

Carrier-derived cell lines and tissues are a precious resource to understand XDP pathology. For example, female heterozygous XDP tissues that are mosaic for functionally wild-type or mutant cells can reveal cell intrinsic *vs* cell extrinsic mechanisms. If neurodegeneration is limited to functionally mutant neurons, then a cell intrinsic mechanism is necessary for degeneration. In contrast if mutant and wildtype neurons degenerate similarly but at a reduced rate compared to male XDP, then cell extrinsic mechanisms, such as network activity properties, degree of inflammation and/or secreted toxic proteins, are likely more important for degeneration than cell intrinsic mechanisms.

The limited number of females with severe XDP examined suggests that mosaic XCI might be a protective factor. An extremely skewed (98%:2%) XCI was noted in only a single case with full XDP syndrome (Domingo et al., 2014), and X-chromosome monosomy (45,X/46,XX) was found in a subset of cells of another female patient with XDP(Westenberger et al., 2013). A recent study examining the peripheral blood of 17 carrier females (4 affected with parkinsonism and 13 non affected) (Al Ali et al., 2021), is also consistent with a mosaic XCI status being protective. Decreased full length TAF1 expression, as measured by the ratio of TAF1 3’/5’, is at an intermediate level for asymptomatic carriers such that XDP patients demonstrated the most reduction and asymptomatic female carriers demonstrated intermediate reduction compared to healthy controls.

Here, for the first time, we reported a sex-differential expression of the *TAF1* gene in stem cell models. We found that XDP female carrier iPSCs as well as a large cohort of female wild-type iPSCs displayed significantly reduced levels of *TAF1* gene, in comparison to male control iPSCs. In our present study, we did not investigate whether there is a correlation between TAF1 changes at mRNA and protein levels in female cells. However, a decreased protein expression, in accordance with reduced TAF1 mRNA expression, was detected in both XDP iPSCs and NSCs, compared to control samples, although the decrease was not statistically significant (Aneichyk et al., 2018; Rakovic et al., 2018).

Decreased expression of X-linked genes is not usually observed in females (Disteche, 2012). Additional studies are needed to determine the downstream effects of sex-linked variation of TAF1 gene expression in iPSCs, particularly in light of its crucial involvement in transcription initiation, and in the regulation of cellular processes, such cell growth (Sekiguchi et al., 1991) or progression through G1 phase of cell cycle (O’Brien and Tjian, 1998).

### TAF1 expression variation in human postmortem brains and XDP

The specific role of TAF1 in the striatum and how it relates to regional differences in pathology has not yet been defined (Bragg et al., 2019). TAF1 is required for postnatal maintenance of striatal cholinergic interneurons in mice (Cirnaru et al., 2021). In humans, rare loss of function mutations in TAF1 cause intellectual disability (Hurst et al., 2018; O’Rawe et al., 2015). Importantly, XDP patients samples have a moderate decrease of TAF1 (∼29% decrease from controls) (Al Ali et al., 2021) which is less severe than the loss of function mutations.

A number of reports have demonstrated decreased TAF1 exon expression downstream of the SVA insertion, both in patient samples and in *in vitro* cellular models (Al Ali et al., 2021; Aneichyk et al., 2018; Domingo et al., 2016; Ito et al., 2016; Makino et al., 2007). However, there are inconsistent reports about which TAF1 transcripts are mainly altered in XDP: the most prominent TAF1 isoform, cTAF1, or the neuron-specific nTAF1 which varies from cTAF1 by only an additional 6 bp derived from the alternative exon 34’ (Capponi et al., 2021; Ito et al., 2016; Makino et al., 2007).

In this study, we have interrogated TAF1 expression variations in both stem cell models and human post-mortem brains. Gene-level quantification, which sums the quantification of all *TAF1* transcripts and does not distinguish transcripts or isoforms, was used for all RNAseq analysis. For analysis of GTEX and Brainseq consortium datasets of postmortem brain, we further investigated differential expression at the individual exon level and found that all exons throughout *TAF1* with sufficient expression for rigorous statistical analysis (n=25) followed the same trend of expression as the gene level quantification. Lower sequencing depth of the iPS samples and 13 exons in the postmortem samples prohibited exon level analysis.

Our data also provided strong evidence that *TAF1* expression varies significantly: between sexes, among different brain regions, and across the life span in humans. The decrease in females is generally modest and of unknown significance, but it is an important consideration when comparing male and female subjects. Of particular relevance to XDP, we found that *TAF1* expression decreased after age 15, specifically in the caudate nucleus. It is important to note that all transcriptome analyses were performed on publicly available datasets generated from bulk tissues, composed of heterogeneous cell types, which do not allow to investigate any differential expression at cellular level.

To our knowledge, there are no other reports of differential age-associated *TAF1* and SVA expression in human postmortem brain tissues. Previously, expression of *TAF1* throughout development and aging was only investigated in mouse brains using quantitative RT-PCR analysis(Jambaldorj et al., 2012). In that study, the authors reported the highest levels of *Taf1* at the embryonic stage, with a gradual reduction till three postnatal weeks, followed by a stable expression of *Taf1* transcript till forty postnatal weeks. While other brain regions including the cerebellum are affected in XDP, the analysis here are restricted to the publicly available human brain lifespan postmortem datasets of GTEX and Brainseq, which do not include early life and adolescent periods in other regions. Therefore, future work will be required to evaluate if TAF1 expression decreases during adolescence in other vulnerable tissue.

### SVA-derived transcripts and XDP

In addition, abundant evidence shows that SVA-driven pathological events lead to XDP dysregulation, although the precise mechanisms underlying the pathophysiology remains elusive. It is proposed that SVA alters local chromatin structure and forms G-quadruplex DNA (G4) (Hancks and Kazazian, 2010), leading to transcriptional interference impeding the progression of RNA polymerase II in XDP samples(Bragg et al., 2017). Also, the SVA insertion includes a hexameric (CCCTCT)_n_ repeat of variable length, which is inversely correlated with age of disease onset in individuals with XDP (Bragg et al., 2017; Westenberger et al., 2019). These findings directly correlate genetic variation in individuals with XDP to differences in their manifestation of clinical disease, thereby supporting a causal role for the SVA insertion in disease pathogenesis.

Therefore, we hypothesized that the regulation of SVA-F expression in the brain may also influence the XDP manifestations. Our data showed slight but significantly decreased SVA expression after 30 years of life in the DLPC and caudate nucleus. No significant difference between sexes or other age groups were observed. Approximately 2700 SVA retrotransposon copies are present in the human genome (Hancks and Kazazian, 2010), creating inherited genetic diversity by actively mutagenizing the genome, and contribute the vast majority of GC-rich human-specific tandem repeat expansions in the human genome (Sulovari et al., 2019). While neurotypical brains do not have the patient specific SVA-F insertion in TAF1 intron 32, other SVA-F copies with identical sequences are present in all humans. These identical SVA sequences are regulated by cellular host factors in trans by members of the large family of protein coding Kruppel-associated box zinc finger proteins (KZFP) bind SVA elements to dock KRAB-associated protein 1 (KAP1/TRIM28) (Rowe et al., 2010). KZFP and KAP/TRIM28 are generally lowly expressed in postnatal striatum, for example the caudate, basal ganglia and putamen are amongst the tissue with the lowest expression of TRIM28, ZNF587, ZNF91/93 in GTEx (GTEx Consortium, 2020). The SVA region harbors extensive DNA methylation in XDP patient blood and the level of methylation is reduced in the basal ganglia of the single patient tissue examined thus far (Lüth et al., 2022). Our findings suggest dynamic regulation of SVA elements in the adult caudate nucleus. It is uncertain which factors cause the decreased SVA-F expression after 30 years but this regulation may modify XDP phenotype.

Currently, the hypothesized causal path is that SVA alters TAF1 and the altered TAF1 leads to XDP phenotype (SVA → TAF1 → pathology). It is possible that XDP specific SVA-derived transcripts directly affect TAF1 and/or that SVA derived transcripts directly affect pathology (SVA → TAF1). Due to the multicopy nature of SVA-F, no reports have conclusively detected XDP specific SVA-derived transcripts. Here, we specifically used stranded total RNA sequencing to detect any XDP specific transcripts. We find weak evidence for the presence of any XDP specific transcripts. Therefore, we conclude SVA-derived transcripts are either not expressed or below the limits of detection.

## Conclusions

We unveiled sex-, tissue- and age-specific regulation of *TAF1* in human brain. We found that, uniquely in the caudate nucleus, where *TAF1* expression is not sexually dimorphic, *TAF1* expression significantly decreases after adolescence. Although these findings were obtained analyzing brain tissues from neurotypical individuals, we hypothesize that the decrease of *TAF1* after 15 years of age may have implications for the XDP adult neostriatum. In our proposed model, the decrease in *TAF1* after adolescence in the caudate nucleus may synergize with the XDP-specific partial loss of *TAF1* function, causing an even further loss of *TAF1*, and thereby triggering degeneration in adult life. This model is consistent with the late onset of XDP neurodegeneration, with patients appearing neurologically unaffected until 35-40 years of age, and with the female carriers only showing attenuated symptoms due to their mosaic neostriatum.

To better frame the role of *TAF1* in the XDP neurodegeneration, and to inform about the causal mechanisms leading to XDP pathology, it is crucial to perform extensive transcriptomic analysis on XDP post-mortem brain tissues. To capture novel cellular phenotypes and mechanisms underlying the striatal degeneration, further studies are needed to differentiate the isogenic XDP female carrier iPSCs reported here, together with patient-derived iPSCs into specific neuronal models. Such studies will help us better understand the impact of *TAF1* loss in iPSC-derived striatal models, as well as the direct or indirect contribution of SVA-driven pathological mechanisms.

## 13. Acknowledgements

We thank Dr. Ronald McKay and Dr. Mike McConnell (Lieber Institute for Brain Development) for their thoughtful feedback throughout all aspects of the study. We are grateful for the contributions of the Office of the Chief Medical Examiner of the District of Columbia; the Office of the Chief Medical Examiner for Northern Virginia, Fairfax Virginia; and the Office of the Chief Medical Examiner of the State of Maryland, Baltimore, Maryland. We also thank the Medical University of Sofia Bulgaria, and Dr. H. Ronald Zielke, Robert D. Vigorito, and Robert M. Johnson of the National Institute of Child Health and Human Development Brain and Tissue Bank for Developmental Disorders, at the University of Maryland, for providing child and adolescent brain specimens. Finally, we are indebted to the generosity of the families of the decedents, who donated the brain tissue used in these studies.

## 14. Conflict of Interest

Authors report no conflict of interest

## 15. Funding sources

This work was supported by the Lieber Institute for Brain Development, the MGH Collaborative Center for X-Linked Dystonia-Parkinsonism (JAE, ACMP, DCB), the Maryland Stem Cell Research Fund (LD), and the National Institutes of Health (KJMB - T32MH015330).

---

**Extended Data Figure 1-1**. **Detection of SVA retrotransposon insertion in the genomic DNA of XDP female carrier-derived iPSC lines. a.** Family trees showing that the iPSC clones were derived by daughters/obligated carriers of XDP probands. **b.** Long-PCR products obtained from amplification of genomic DNA extracted from one control, one XDP mutant line, and all eight XDP female carrier-derived iPSC clones. Primers flanking the SVA insertion site were used to confirm the presence of the SVA retrotransposon in the genome. The control iPSC line shows only a product of ∼599 bp (lower band), whereas the male XDP-derived iPSC line shows the predicted 3229 bp SVA product (upper band). Both PCR products were detectable in all eight XDP female carrier-derived iPSCs.

*Figure contribution*: Laura D’Ignazio performed the experiment and analyzed data.

**Extended Data Figure 1-2. Pluripotency characterization of XDP female carrier-derived iPSCs. a.** Bulk RNA-sequencing analysis revealed that all XDP female carrier-derived iPSC clones expressed multiple markers associated with pluripotency, such as *SOX2*, *POU5F1* and *NANOG*; they did not express genes associated with three germ layers: *EOMES* (mesoderm), *GATA4* (endoderm) and *PAX6* (neuroectoderm). **b.** Expression of the pluripotency markers NANOG (green) and TRA-1-60 (red) was detected by immunofluorescence staining in all eight XDP carrier iPSCs. Hoechst (blue) was used to visualize nuclei. Merged images depict overlays of immunoreactivity for each target, together with the nuclear counterstain. Scale bar = 20 μm **c.** Comparative expression of primed (*DUSP6, DNMT3A, SOX11*) and naive (*DPPA5, KLF17, DNMT3L*) markers in all eight XDP female carrier-derived iPSCs. **d.** Representative images from isogenic XDP female carrier iPSC lines showing localization of RNA scope-specific probes for DUSP6 (green) transcripts. DAPI was used to stain nuclei. Scale bar = 20 µm.

*Figure contribution*: Bareera Qamar performed the fluorescence immunostaining on iPSCs; Laura D’Ignazio performed the RNA scope assay; Kynon J.M. Benjamin performed the RNA-seq data analysis.

**Extended Data Figure 1-3. XCI status of XDP female carrier-derived iPSCs. a.** Principal component analysis of normalized expression of X-linked genes from the eight XDP carrier iPSC lines. **b.** Bar plots of the percentage of allele-specific transcripts derived from chromosome 7, which are shared between the indicated iPSC clones. Chromosome 7 was chosen as a reference autosomal chromosome because it has a similar size to chromosome X. Expression from the same or different alleles was determined by counting the number of shared and unique homozygous single nucleotide polymorphisms (SNPs), respectively. The number of SNPs analyzed for each pair is shown at the right of each bar. **c.** Diagram illustrating differences between “proper” and “eroded” XCI status. Proper XCI occurs when X-linked genes show higher monoallelic expression than do autosomal genes. Erosion of XCI occurs when the fraction of monoallelic transcripts is similar between the X chromosome and autosomes. **d.** Fraction of monoallelic expression from chromosome X compared to that from all autosomal chromosomes. An increased fraction of monoallelic expression indicates silencing of the X chromosome, and that proper XCI occurs in all XDP female carrier-derived iPSCs. **e.** Summary of the XCI status of each iPSC clone.

*Figure contribution*: Kynon J.M. Benjamin performed the PCA analysis; Ricardo S. Jacomini and Apua C.M. Paquola performed the allele-specific transcriptomic analysis.

**Extended Data Figure 3-1. *XIST* and *XACT* nuclear localization in XDP carrier-derived isogenic iPSCs. a.** Representative images showing localization of *XIST*(red) and *XACT* (green) RNA scope probes in 33360.A and 33360.B iPSCs. DAPI was used to stain nuclei. Scale bar = 20 µm. **b.** Percentage of XIST-positive cells where *XIST* and *XACT* lncRNAs have distinct or shared nuclear localization in the set of isogenic XDP female carrier-derived iPSCs. For 33360.B, iPSCs also show the percentage of *XIST*+ cells not expressing *XACT*.

*Figure contribution*: Bareera Qamar and Laura D’Ignazio performed the RNA scope assay and analyzed data.

**Extended Data Figure 5-1. Sex differential expression analysis of *TAF1* and *XIST* across regions of postmortem adult brains. a.** Box plots comparing residualized gene expression of TAF1, in female (red) and male (blue) individuals, from the BrainSeq Consortium (Benjamin et al., 2020; Collado-Torres et al., 2019) across the dorsolateral prefrontal cortex (DLPFC; n = 379) and hippocampus (n = 376). ns = not significant. **b.** Box plots comparing residualized gene expression of XIST in female (red) and male (blue) individuals from the BrainSeq Consortium (Benjamin et al., 2020; Collado-Torres et al., 2019); caudate nucleus (n = 394), dorsolateral prefrontal cortex (DLPFC; n = 379), and hippocampus (n = 376). **** p ≤ 0.001.

*Figure contribution*: Ria Arora and Kynon J.M. Benjamin performed the sex differential expression analysis.

**Extended Data Figure 5-2. SVA expression analysis in IPS lines and postmortem caudate nucleus. a.** Box plots comparing residualized gene expression of subfamily F of the SVA retrotransposon in the caudate nucleus (n = 394), DLPFC (n=379), and hippocampus (n=376) of female (red) and male (blue) individuals from the BrainSeq Consortium (Benjamin et al., 2020; Collado-Torres et al., 2019). ns = not significant. **b-c.** Box plots comparing residualized gene expression in individuals from the BrainSeq Consortium (Benjamin et al., 2020; Collado-Torres et al., 2019) for **(b)** SVA-F in the dorsolateral prefrontal cortex (DLPFC), and **(c)** SVA-F in the hippocampus. Samples were divided into four groups based on the age of the individuals: 0-15 years; 16-31 years; 31-51 years; and older than 51 years. ns = not significant; **FDR ≤ 0.01. **d.** SVA expression in iPS cells. Plot shows normalized RNA-Seq coverage along the SVA sequence for both sense (+) and antisense (-) strands. Mutant trace (red) indicates female XDP carrier lines expressing the XDP allele: 33110.2A, 33110.2E, 33110.2F, 33360.A, 33360.D; Wild-type trace (light blue) indicates female XDP carrier lines expressing the wild-type X chromosome: 33360.B, 33811.A, 33811.B. The schematic illustration describes the structure of the human SVA retrotransposons.

*Figure contribution*: Taylor A. Evans performed the SVA-F differential expression analyses; Apua C.M. Paquola and Jennifer A. Erwin performed the SVA-derived transcript analysis on iPSCs.

**Extended Data Table 1-1**. Characteristics of 11 human iPSCs used in this study, including genotype, clone, genotype of clone, number of SVA hexameric repeats, source and PMID. ΔSVA-XDP denotes isogenic SVA-deleted XDP. For the XDP proband (33109 male, clone 2B) age at onset was known, 58 years.

**Extended Data Table 5-1.** Genes and repeats significantly downregulated, with FDR < 0.05, in the caudate nucleus over life span. Samples from neurotypical control individuals from the BrainSeq Consortium (Benjamin et al., 2020; Collado-Torres et al., 2019) were divided in four age groups: A) 0-15 years; B) 16-30 years; C) 31-51 years; D) older than 51 years. Three pairwise comparisons were performed among age groups (A vs B, A vs C, and A vs D).

**Extended Data Table 5-2**. Differential residualized expression of TAF1 exons over the life span. Caudate nucleus, DLPFC and hippocampus samples from neurotypical control individuals from the BrainSeq Consortium (Benjamin et al., 2020; Collado-Torres et al., 2019) were divided into five age groups: PRE) prenatal; A) 0-15 years; B) 16-30 years; C) 31-51 years; D) older than 51 years. Ten pairwise comparisons were performed among age groups for every TAF1 exon passing a low expression filter (PRE vs A, PRE vs B, PRE vs C; PRE vs D; A vs B, A vs C, A vs D; B vs C, B vs D; C vs D).

## Supplementary Figure Legends and Table Titles

**Supplementary Fig. S1.**
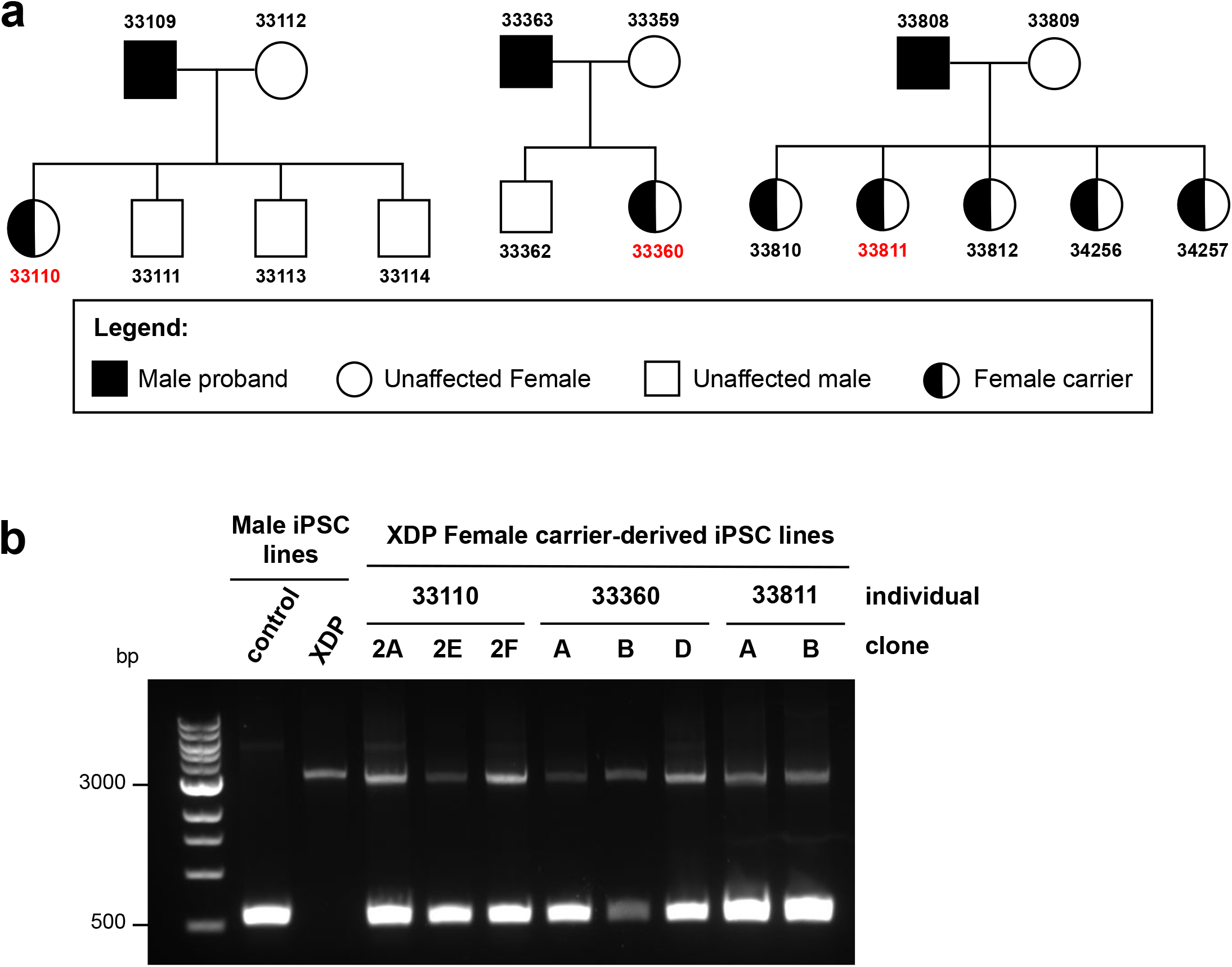
Detection of SVA retrotransposon insertion in the genomic DNA of XDP female carrier-derived iPSC lines. **a.** Family trees showing that the iPSC clones were derived by daughters/obligated carriers of XDP probands. **b.** Long-PCR products obtained from amplification of genomic DNA extracted from one control, one XDP mutant line, and all eight XDP female carrier-derived iPSC clones. Primers flanking the SVA insertion site were used to confirm the presence of the SVA retrotransposon in the genome. The control iPSC line shows only a product of ∼599 bp (lower band), whereas the male XDP-derived iPSC line shows the predicted 3229 bp SVA product (upper band). Both PCR products were detectable in all eight XDP female carrier-derived iPSCs.

**Supplementary Fig. S2.**
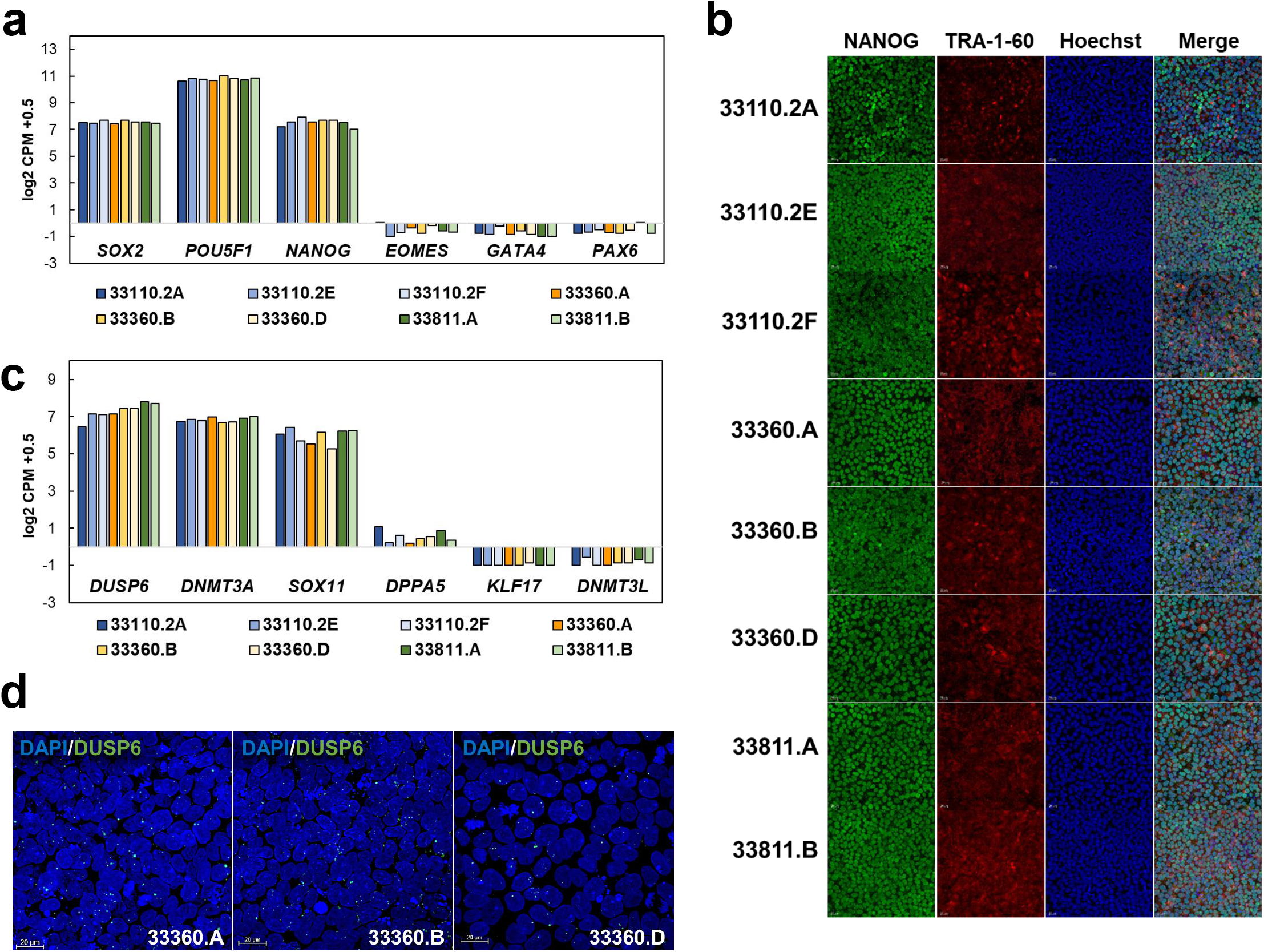
Pluripotency characterization of XDP female carrier-derived iPSCs. **a.** Bulk RNA-sequencing analysis revealed that all XDP female carrier-derived iPSC clones expressed multiple markers associated with pluripotency, such as *SOX2*, *POU5F1* and *NANOG*; they did not express genes associated with three germ layers: *EOMES* (mesoderm), *GATA4* (endoderm) and *PAX6* (neuroectoderm). **b.** Expression of the pluripotency markers NANOG (green) and TRA-1-60 (red) was detected by immunofluorescence staining in all eight XDP carrier iPSCs. Hoechst (blue) was used to visualize nuclei. Merged images depict overlays of immunoreactivity for each target, together with the nuclear counterstain. Scale bar = 20 μm **c.** Comparative expression of primed (*DUSP6, DNMT3A, SOX11*) and naive (*DPPA5, KLF17, DNMT3L*) markers in all eight XDP female carrier-derived iPSCs. **d.** Representative images from isogenic XDP female carrier iPSC lines showing localization of RNA scope-specific probes for DUSP6 (green) transcripts. DAPI was used to stain nuclei. Scale bar = 20 µm.

**Supplementary Fig. S3.**
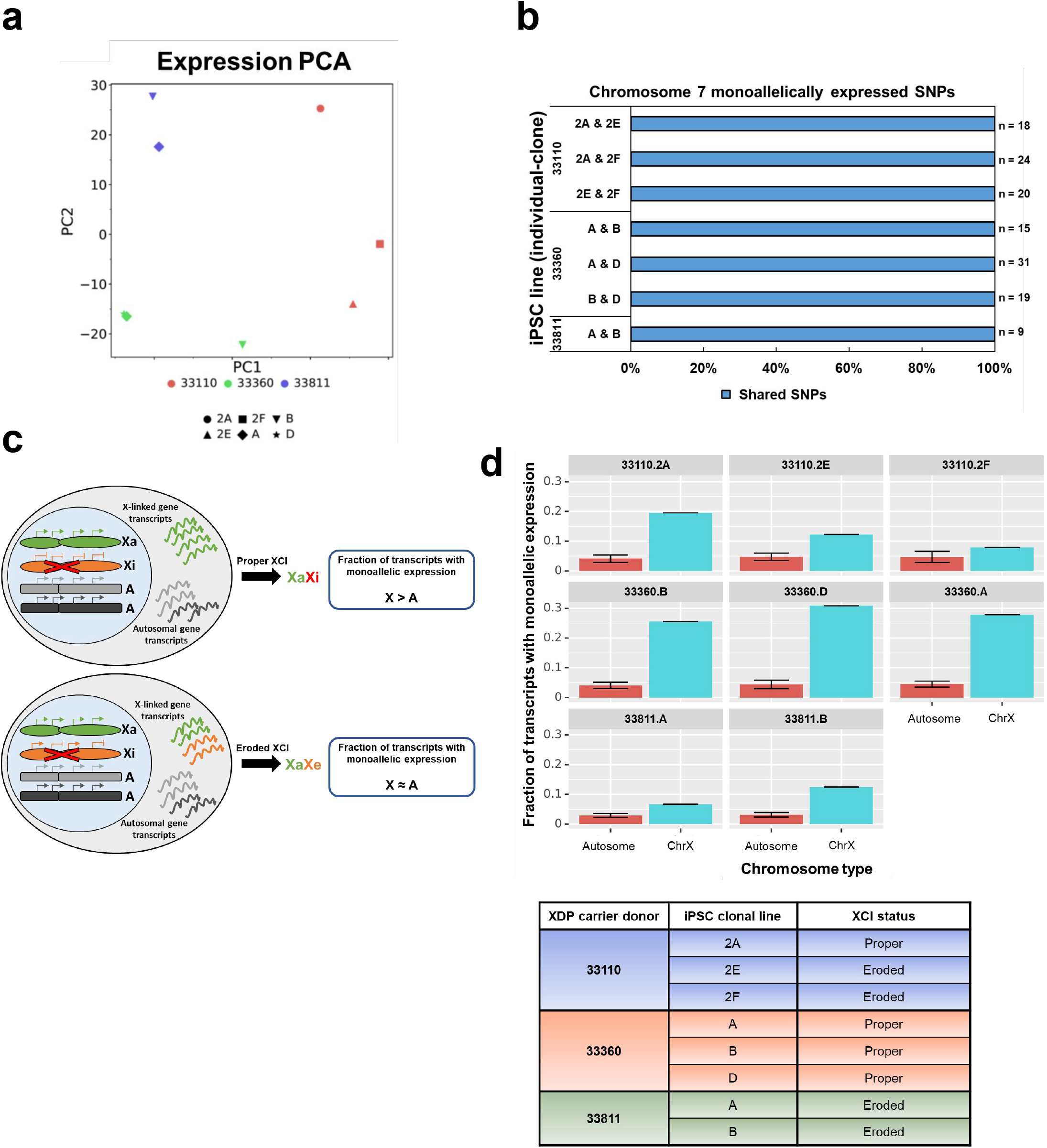
XCI status of XDP female carrier-derived iPSCs. **a.** Principal component analysis of normalized expression of X-linked genes from the eight XDP carrier iPSC lines. **b.** Bar plots of the percentage of allele-specific transcripts derived from chromosome 7, which are shared between the indicated iPSC clones. Chromosome 7 was chosen as a reference autosomal chromosome because it has a similar size to chromosome X. Expression from the same or different alleles was determined by counting the number of shared and unique homozygous single nucleotide polymorphisms (SNPs), respectively. The number of SNPs analyzed for each pair is shown at the right of each bar. **c.** Diagram illustrating differences between “proper” and “eroded” XCI status. Proper XCI occurs when X-linked genes show higher monoallelic expression than do autosomal genes. Erosion of XCI occurs when the fraction of monoallelic transcripts is similar between the X chromosome and autosomes. **d.** Fraction of monoallelic expression from chromosome X compared to that from all autosomal chromosomes. An increased fraction of monoallelic expression indicates silencing of the X chromosome, and that proper XCI occurs in all XDP female carrier-derived iPSCs. **e.** Summary of the XCI status of each iPSC clone.

**Supplementary Fig. S4.**
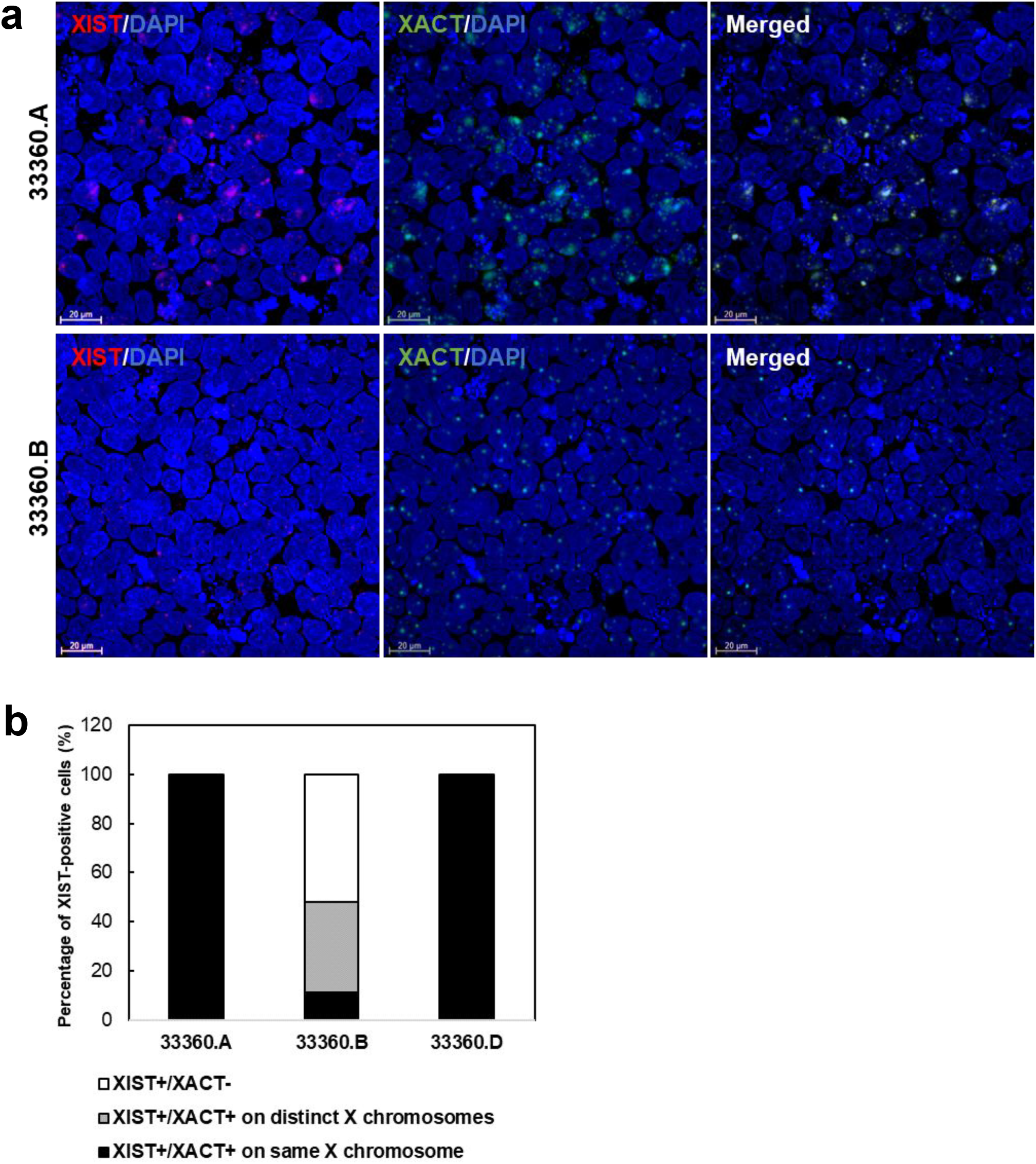
*XIST* and *XACT* nuclear localization in XDP carrier-derived isogenic iPSCs. **a.** Representative images showing localization of *XIST* (red) and *XACT* (green) RNA scope probes in 33360.A and 33360.B iPSCs. DAPI was used to stain nuclei. Scale bar = 20 µm. **b.** Percentage of XIST-positive cells showing localization of *XIST* and *XACT* lncRNAs on the same, or different X chromosomes, in the set of isogenic XDP female carrier-derived iPSCs. For 33360.B, iPSCs also show the percentage of *XIST*+ cells not expressing *XACT*.

**Supplementary Fig. S5.**
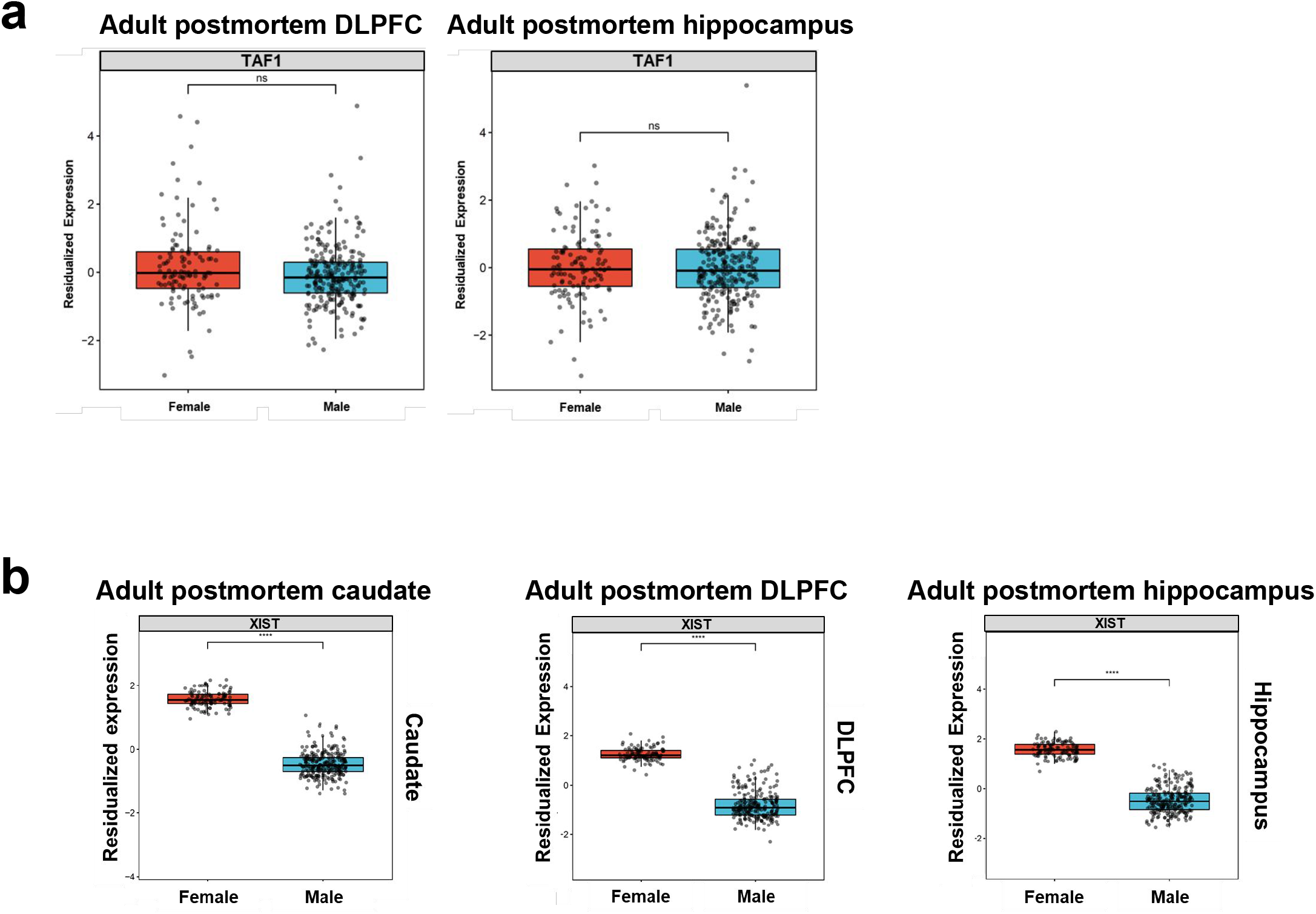
Sex differential expression analysis of *TAF1* and *XIST* across regions of postmortem adult brains. **a.** Box plots comparing residualized gene expression of TAF1, in female (red) and male (blue) individuals, from the BrainSeq Consortium (Benjamin et al., 2020; Collado-Torres et al., 2019) across the dorsolateral prefrontal cortex (DLPFC; n = 379) and hippocampus (n = 376). ns = not significant. **b.** Box plots comparing residualized gene expression of XIST in female (red) and male (blue) individuals from the BrainSeq Consortium (Benjamin et al., 2020; Collado-Torres et al., 2019); caudate nucleus (n = 394), dorsolateral prefrontal cortex (DLPFC; n = 379), and hippocampus (n = 376). **** p ≤ 0.001.

**Supplementary Fig. S6.**
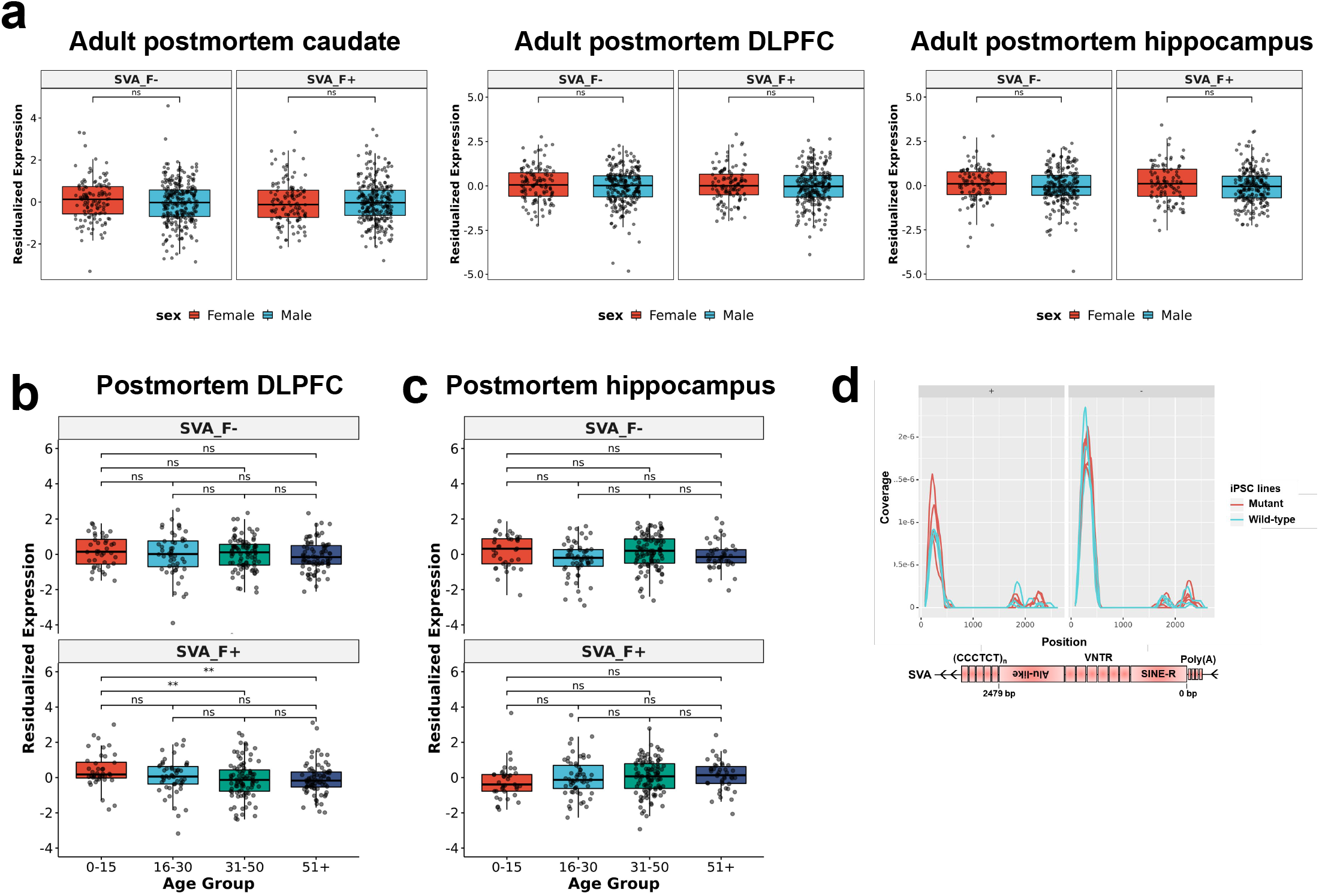
SVA expression analysis in IPS lines and postmortem caudate nucleus. **a.** Box plots comparing residualized gene expression of subfamily F of the SVA retrotransposon in the caudate nucleus (n = 394), DLPFC (n=379), and hippocampus (n=376) of female (red) and male (blue) individuals from the BrainSeq Consortium (Collado-Torres et al. 2019; Benjamin et al. 2020). ns = not significant. **b-c.** Box plots comparing residualized gene expression in individuals from the BrainSeq Consortium (Benjamin et al., 2020; Collado-Torres et al., 2019) for **(b)** SVA-F in the dorsolateral prefrontal cortex (DLPFC), and **(c)** SVA-F in the hippocampus. Samples were divided into four groups based on the age of the individuals: 0-15 years; 16-31 years; 31-51 years; and older than 51 years. ns = not significant; **FDR ≤ 0.01. **d.** SVA expression in iPS cells. Plot shows normalized RNA-Seq coverage along the SVA sequence for both sense (+) and antisense (-) strands. Mutant trace (red) indicates female XDP carrier lines expressing the XDP allele: 33110.2A, 33110.2E, 33110.2F, 33360.A, 33360.D; Wild-type trace (light blue) indicates female XDP carrier lines expressing the wild-type X chromosome: 33360.B, 33811.A, 33811.B. The schematic illustration describes the structure of the human SVA retrotransposons.

**Supplementary Table S1.** Characteristics of 11 human iPSCs used in this study, including genotype, clone, genotype of clone, number of SVA hexameric repeats, source and PMID. ΔSVA-XDP denotes isogenic SVA-deleted XDP. For the XDP proband (33109 male, clone 2B) age at onset was known, 58 years.

**Supplementary Table S2.** Genes and repeats significantly downregulated, with FDR < 0.05, in the caudate nucleus over life span. Samples from neurotypical control individuals from the BrainSeq Consortium (Benjamin et al., 2020; Collado-Torres et al., 2019) were divided in four age groups: A) 0-15 years; B) 16-30 years; C) 31-51 years; D) older than 51 years. Three pairwise comparisons were performed among age groups (A vs B, A vs C, and A vs D).

**Supplementary Table S3.** Differential residualized expression of TAF1 exons over the life span. Caudate nucleus, DLPFC and hippocampus samples from neurotypical control individuals from the BrainSeq Consortium (Benjamin et al., 2020; Collado-Torres et al., 2019) were divided into five age groups: PRE) prenatal; A) 0-15 years; B) 16-30 years; C) 31-51 years; D) older than 51 years. Ten pairwise comparisons were performed among age groups for every TAF1 exon passing a low expression filter (PRE vs A, PRE vs B, PRE vs C; PRE vs D; A vs B, A vs C, A vs D; B vs C, B vs D; C vs D).

## Notes

### Competing Interest Statement

The authors have declared no competing interest.

